# The CTEXT complex in *Saccharomyces cerevisiae* plays a crucial role in degrading distinct sets of aberrant mRNAs by the nuclear exosome

**DOI:** 10.1101/2021.03.29.437469

**Authors:** Upasana Saha, Rajlaxmi Gaine, Sunirmal Paira, Satarupa Das, Biswadip Das

## Abstract

In *Saccharomyces cerevisiae*, DRN (**<u>D</u>**ecay of **<u>R</u>**NA in the **<u>N</u>**ucleus) requiring Cbc1/2p, Tif4631p, and Upf3p promotes the exosomal degradation of aberrantly long 3′-extended-, export-defective transcripts and a small group of normal (special) mRNAs. In this study, using a systematic proteomic analysis we show that each of the known components interacts with one another and they exist as a separate complex, which was dubbed CTEXT (**<u>C</u>**BC-**<u>T</u>**if4631p-dependent **<u>EX</u>**osome **<u>T</u>**argeting). We also identified a DEAD-box RNA helicase Dbp2p as an additional novel component of CTEXT during this analysis which was further bolstered by the finding that genomic deletions of Dbp2p led to the stabilization of all the signature nuclear messages. Interestingly, the RRM domain of Tif4631p located at the extreme N-termini of this polypeptide was found to play a vital role in in mediating the interaction of the CTEXT with the core exosome complex. These inferences were substantiated by the finding that deletion of this domain led to the functional impairment of the CTEXT complex. Thus, the CTEXT constitutes an independent complex that assists the nuclear exosome in degrading the select classes of nuclear transcripts in *Saccharomyces cerevisiae*.

## Introduction

In *Saccharomyces cerevisiae* an elaborate array of mRNA surveillance and quality control mechanisms eliminate aberrant messages generated from the error-prone nuclear mRNP biogenesis (Parker 2012; Das and Das 2013; Mitchell 2014; Kilchert et al. 2016). The nuclear exosome targets and degrades a wide array of aberrant and normal mRNAs (Mitchell et al. 1997; Allmang et al. 1999a, b, 2000; Butler 2002; Parker 2012; Das and Das 2013; Singh et al. 2018). In the nucleus, the exosome targets transcription-elongation-defective (e.g., pre-rRNAs, pre-sno RNAs and pre-tRNAs) (Gudipati et al. 2012; Schneider et al. 2012) intron-containing splice defective pre-mRNAs (Bousquet-Antonelli et al. 2000; Gudipati et al. 2012; Kong et al. 2014), aberrantly long 3′-extended mRNAs (Das et al. 2000; Torchet et al. 2002; Maity et al. 2016) and export defective faulty nuclear transcripts (Das et al. 2003, 2006; Kuai et al. 2005; Maity et al. 2016). The nuclear exosome consists of nine catalytically inactive core subunits, two associated processive subunits and a few nuclear cofactors (Butler 2002; Liu et al. 2006; Parker 2012; Wasmuth et al. 2014). Nine core subunits are arranged in the form of a barrel (known as EXO9), which are arranged in two stacked ring structures. The top ‘trimeric’ cap (composed of Rrp4p Rrp40p and Csl4p) is placed on the top of a bottom ‘hexameric’ ring (consists of Rrp41p, Rrp42p, Rrp43, Rrp45p, Rrp46p and Mtr3p) with a central channel. EXO9 contacts two catalytically active tenth and eleventh subunits, Dis3p/Rrp44p (possess both an endo-and a 3′→5′ exoribonuclease activity) and Rrp6p (3′→ 5′ exoribonuclease), respectively, from opposite sides to form EXO11^Dis3p+Rrp6p^ (Dziembowski et al. 2007; Makino et al. 2013a, b; Wasmuth et al. 2014).

The selectivity of the nuclear exosome to target diverse RNA substrates requires assistance from exosomal co-factors, which aids the exosome to specifically recognize a given kind of RNA target as well as modulates the catalytic activity of the exosome (Allmang et al. 2000; van Hoof et al. 2000; Kadaba et al. 2004; Wlotzka et al. 2011; Schmidt and Butler 2013). Functionally, these co-factors are thought to primarily recognize a specific RNA target and then further recruit them on to EXO11^Dis3p+Rrp6p^ to facilitate their degradation (Schmidt and Butler 2013). TRAMP (**<u>TR</u>**f4p/5p-**<u>A</u>**ir1p/2p-**<u>M</u>**tr4p-**<u>P</u>**olyadenylation) complex in *S. cerevisiae* provides the best understood and well-studied example of such ancillary complex, which consists of DExH box RNA helicase, Mtr4p (Halbach et al. 2013; Hardwick and Luisi 2013; Kilchert et al. 2016), non-canonical poly(A) polymerase, Trf4p (Pap2p)/Trf5p and Zn-knuckle RNA binding proteins, Air1p/2p (Hardwick and Luisi 2013; Kilchert et al. 2016). In addition, another nuclear mRNA degradation system dubbed DRN (**<u>D</u>**ecay of the m**<u>R</u>**NA in the **<u>N</u>**ucleus) that relies on the nuclear-cap binding protein Cbc1p and Rrp6p was independently identified, which was known to degrade aberrant *cyc1-512* (Das et al. 2000) and *lys2-187* mutant mRNAs (Das et al. 2006) and a small subset of normal mRNAs dubbed special mRNAs (Kuai et al. 2005). Further genetic analysis showed that DRN system requires the product of a few additional genes including Cbc2p, Tif4631p, and Upf3p (Das et al. 2014) based on the finding that mutation/deletion of any one of these genes led to the stabilization and diminished decay of the DRN substrate mRNAs, mutants *cyc1-512*, *lys2-187* mRNAs and select special mRNAs (Das et al. 2014). Furthermore, mutations/deletions in these genes displayed a strong epistasis among themselves as well as with the mutations in the components of the core nuclear exosome (Das et al. 2014; Maity et al. 2016) with respect to the stabilization and nuclear decay. These findings suggest that together Cbc1p/2p, Tif4631p, Upf3p and Rrp6p are playing a role in the degradation of *cyc1-512, lys2-187* and select special mRNAs. However, the functional relationship of DRN with the nuclear exosome and its cofactor TRAMP remained undefined.

A functional relationship between DRN and the nuclear exosome/TRAMP was established by demonstrating that DRN system appear to serve as a cofactor of the nuclear exosome (Maity et al. 2016). Our analysis initially categorized the known spectra of the aberrant nuclear mRNAs in *Saccharomyces cerevisiae* into four broad classes (Maity et al. 2016) including (i) transcription elongation defective (Libri et al. 2002; Strasser et al. 2002; Zenklusen et al. 2002; Rougemaille et al. 2007; Assenholt et al. 2008), (ii) intron-containing splice-defective (Bousquet-Antonelli et al. 2000; Gudipati et al. 2012; Schneider et al. 2012; Kong et al. 2014), (iii) aberrantly long 3′-extended transcripts (Birse et al. 1998; Minvielle-Sebastia and Keller 1999; Das et al. 2000; Yonaha and Proudfoot 2000; Torchet et al. 2002; Proudfoot 2016), and export defective mRNAs (Das et al. 2003, 2006; Kuai et al. 2005). We provided evidence of an intriguing effects of core nuclear exosome, TRAMP and DRN on the stability and decay rates of these mRNAs. While the surveillance/degradation of the transcription-elongation- and splice-defective mRNAs require the involvement of the core nuclear exosome and TRAMP, the export defective messages require the participation of the core exosome and DRN. Remarkably, the nuclear surveillance of the aberrant message with abnormally long 3′-extension demands the participation of the core nuclear exosome, TRAMP and DRN systems (Maity et al. 2016). These observations led us to conclude that DRN system, just like TRAMP complex assists the nuclear exosome to degrade a distinct class of aberrant mRNAs either directly (export defective messages) or along with the TRAMP complex (aberrant transcripts with abnormally long 3′-extension) thereby acting as an alternative co-factor of nuclear exosome (Maity et al. 2016). Furthermore, DRN along with the nuclear exosome were shown to regulate multiple cellular processes like Unfolded Protein Response (Sarkar et al. 2018), ability of yeast cells to adapt to low nutrient levels (Das et al. 2019) as well as the maintenance of telomere length in yeast (M. Banerjea and B. Das, unpublished). Thus, DRN along with the nuclear exosome appears to play a pivotal role in regulating vital cellular processes by targeting and preferentially degrading select normal mRNAs in addition to its role in limiting aberrant messages.

Although DRN evolved as an alternative co-factor of the nuclear exosome (Maity et al. 2016) since its initial description as a Cbc1p-dependent nuclear mRNA degradation system (Das et al. 2000), a compelling biochemical evidence whether the components of DRN display physical interactions *in vivo* among themselves or with the components of the core exosome/TRAMP is currently lacking. Moreover, neither an account of a comprehensive composition of the DRN proteome is available, nor it is known if these components exist as a separate physical entity. Strikingly, one of the major DRN component Tif4631p was previously hypothesized to act as an integrator of various protein complexes playing pivotal roles in the distinct aspects of RNA metabolism (Das and Das 2016), such as splicing of a small group of pre-mRNAs (Kafasla et al. 2009) and initiation of translation (Goyer et al. 1993; Tarun Jr. and Sachs 1996, 1997; Wells et al. 1998). Therefore, in this paper we made an effort to dissect out the functional feature of Tif4631p, which revealed that the N-terminal RRM1 domain of Tif4631p plays a vital role in the nuclear mRNA surveillance. To further gain an insight into the exact role of Tif4631p in the functional contribution of the DRN/exosome, we considered if this protein plays any fundamental role in (i) the complex formation of the other components of DRN and (ii) mediating the interaction if DRN with exosome. We used Tif4631p as a bait to co-purify the Tif4631p-interacting proteins by employing Tandem Affinity Purification (TAP) coupled to LC-MS and co-immuno-precipitation. Our analysis (i) provides evidence that Cbc1p, Tif4631p, Dbp2p and Upf3p exists as a separate complex, (ii) identifies Dbp2, a ATP-dependent DEAD-box RNA helicase, that defines a novel component of DRN, and (iii) shows that the four protein complex together displays a physical interaction with the core exosome complex that is mediated via the RRM1 domain of Tif4631p. We dub this integral protein complex consisting of Cbc1p-Tif4631p-Upf3p-Dbp2p **<u>CTEXT</u>** (**<u>C</u>**bc1p-**<u>T</u>**if4631p-dependent-**<u>EX</u>**osomal **<u>T</u>**argeting) complex that together act as an exosomal co-factor that helps exosome to target a distinct sets of aberrant and special nuclear mRNA targets.

## Materials and methods

### Nomenclature, strains, media, and yeast genetics

Standard genetic nomenclature is used to designate wild-type alleles (e.g., *CYC1*, *LYS2*, *NUP116*, and *TIF4631*), recessive mutant alleles (e.g., *cyc1-512*, *lys2-187*, *nup116*-Δ, *tif4631-1*) and disruptants or deletions (e.g., *cbc1*::*URA3*, *cbc1-*Δ). An example of denoting a protein encoded by a gene is as follows: Tif4631pencoded by *TIF4631*. CBC denotes the cap-binding complex composed of Cbc1p and Cbc2p.The genotypes of *S. cerevisiae* strains used in this study are listed in Supplementary Table S1. Standard YPD, YPG, SC-Leu (leucine omission) and other omission media were used for testing and for growth of yeast (Sherman, 1991). Yeast genetic analysis was carried out by standard procedures (Sherman, 1991).

**Table 1.** Various domain deletion mutants of TIF4631 gene and the features and properties of these mutants

| **Designation** | **Domain which is absent in the mutant** | **Genetic symbol/ representation** | **Length of Deletion (in aa)** | **Span of deletion (in aa)** | **Ability to suppress *LCY* allele** | **Ability to suppress *lys2-187* allele** | **Stability of export defective mRNAs** |
| --- | --- | --- | --- | --- | --- | --- | --- |
| Full length | None | Tif4631p | 0 | None | – | – | – |
| ΔRRM1 | RRM1 | Tif4631p-ΔRRM1 | 76 | 5-80 | + | + | + |
| ΔPAB | PAB1-interacting domain | Tif4631p-ΔPAB1 | 137 | 188-324 | + | + | + |
| ΔCBC | CBC1-interacting domain | Tif4631p-ΔCBC | 179 | 437-615 | – | – | – |
| ΔRRM2 | RRM2 | Tif4631p-ΔRRM2 | 64 | 486-549 | – | – | – |
| ΔMIF4G | MIF4G | Tif4631p-ΔMIF4G | 253 | 607-859 | – | – | – |
| ΔRRM3 | RRM3 | Tif4631p-ΔRRM3 | 81 | 862-942 | – | – | – |

### Extraction and purification of Total RNA

Total RNA was isolated from appropriate strains of *S. cerevisiae* as described by Das *et al*. (2000, 2003). Appropriately grown yeast cells were washed with washing buffer after harvesting at 4°C, followed by extraction of total cellular RNA by vortexing in the presence of phenol-chloroform-IAA (25:24:1) for three times. Following extraction, RNA was purified and recovered by precipitation with RNAase free ethanol and further quantified by UV absorption spectrophotometry.

### Analysis of the steady state levels and stability of specific mRNAs

The stabilities of various mRNAs were determined by inhibition of transcription with the transcription inhibitor thiolutin, as described by Das *et al*. (2000, 2003). Briefly, yeast strains were grown at 30°C till mid-logarithmic phase when thiolutin was added to the growing culture at 4µg/ml final concentration followed by withdrawal of a 25 ml of culture at various times after thiolutin addition. Steady state levels of specific mRNAs were determined by harvesting yeast cells at 30°C. Message levels were determined by either real-time PCR (described below) or by Northern blot analysis as described (Das *et al*., 2000, 2003), except that probe was prepared using the psoralen-biotin labeling kit (Applied Biosystems) and mRNA levels quantified using a phosphor-imager. Signals associated with specific message were normalized against *SCR1* signals. The decay rates and half-lives of specific mRNAs were estimated with the regression analysis program (Origin Pro 8, OriginLab Corporation, MA, USA) using a single exponential decay formula y =100 e^-bx^, assuming mRNA decay follows a first order kinetics.

### RT-PCR, DNase Treatment and cDNA Preparation

For reverse transcription, total RNA was treated with DNase I (Fermentas Inc., Pittsburgh, PA, USA). RNA purity was assessed by capillary electrophoresis (Agilent 2100 Bioanalyzer; Agilent Technologies, Inc., Santa Clara, CA) to determine RNA integrity numbers (range 8.7 to 9.5). DNase I-treated RNA (0.5 μg) was reverse transcribed with random primers using Superscript III enzyme (Invitrogen, Carlsbad, CA) for one hour at 50°C. The resultant cDNA was purified with the QIAquick PCR purification system (Qiagen).RNA and cDNA were quantified using the Quant-iT RNA assay kit and the Quant-iT dsDNA HS assay kit respectively with the Qubit® fluorimeter. (Invitrogen, Carlsbad, CA).

### Real Time PCR

1 ng of quantified cDNA was used to quantify the levels of mRNAs by Q-PCR using target specific primers and standard Cyber Green Technology with Power Cyber® Green PCR Master Mix (Applied Biosystem). Q-PCR assays were run in triplicate and conducted using an Applied Biosystems StepOne™ real time PCR system. For each target, purified DNA templates were prepared by PCR, followed by gel purification and serial dilutions to generate standard curves from 10^2^ to 10^9^copy number/reaction versus threshold cycle (C_t_), determined by the StepOne™ software version 2.2.2 (Applied Biosystems). Standard curves were used to determine the amplification efficiencies (E =10^(−1/n)^, where n = slope of the standard curve) in Cyber Green assays as well as copy numbers of templates in experimental samples.

### Statistical Analysis

For all the experiments reported in this paper, three independent (from independent biological replicates) sample size were chosen in the course of all the experiments (*N*=3). However, in some instances, the sample sizes are four or five. Each experiment was performed at least 3 times in all cases, 4 and 5 times in few cases. In each biological replicate, the given strain is grown and treated under the same conditions independently each time while technical replicate is after initial experiments and techniques the exact same sample is analyzed many times. The point of such a technical replicate would be to establish the variability (experimental error) of the analysis technique, thus allowing one to set the confidence limits for what is significant data. The technical replicates are only to control for variants, they are not biological variants. Values which are varying too much from the rest of the values in the specific group were excluded keeping in mind that the variation could be the result of handling and instrumental errors. Statistical analyses were performed using paired sample test. All the statistical parameters such as mean, standard deviations, P-values, standard error of the mean were estimated using OriginPro 8 SRO, version 8.0724 (OriginLab corporation, Northampton, MA, USA). GraphPad 7 software package (GraphPad) was used for preparing relevant graphs for all experiments.

### Preparation of Cell Lysate from *Saccharomyces cerevisiae*

50ml of YPD medium is seeded with sufficient amount of pre-inoculum and incubated at 37°C with shaking until OD_600_ 2.7-3.0 is reached. The cells are then pelleted, washed and resuspended in 1ml of Buffer A (50mM Tris-HCl pH= 7.5, 150mM NaCl, 5mM EDTA, 1mM DTT, 1mM Pmsf, 0.15% NP-40, 1.5mM MgCl_2_, 1x Protease Inhibitor Cocktail (Abcam Cat No: ab201111)). The cells are lysed by vortexing with 20second pulses using glass beads (Unigenetics Instruments Pvt. Ltd Cat No: 11079105). After cell extraction the mixture is centrifuged to get rid of cell debris and glass beads. The supernatant is collected into a fresh microcentrifuge tube and centrifuged again at 2500rpm at 4°C for 10 minutes to get rid of any carried over cell debris and glass beads. The supernatant is collected again into a fresh microcentrifuge tube and centrifuged again at 15300rpm at 4°C for 45 minutes. The supernatant now obtained is clear cell lysate from *Saccharomyces cerevisiae* ready for downstream processing. The cell lysate is quantified using Coomassie Plus (Bradford) Assay Kit (Thermo Scientific) according to standard Bradford assay protocol (Bradford, M.M, 1976).

### TAP Purification

The cell lysate thus obtained was processed according to the protocol from Seraphin *et. al* (2001). The cell lysate is incubated with 100ul of equilibrated IgG slurry (IgG Sepharose^TM^ 6 Fast Flow from GE Healthcare) overnight in rotator wheel at 4°C with slow rotation. Following this the next day, beads were collected by centrifugation and washed two times with Buffer A at 4°C in rotator wheel. The beads are finally collected by centrifugation and resuspended in required amount of autoclaved distilled water to carry out TEV cleavage with Promega ProTEV Plus (V610A) according to manufacturers protocol. The beads were centrifuged and this time the eluate was collected in a fresh microcentrifuge tube. The eluate thus obtained was incubated with equilibrated 200ul Calmodulin beads (Calmodulin Sepharose^TM^ 4B from GE Healthcare) in Calmodulin Binding Buffer (10mM Tris-HCl pH=7.5,150mM NaCl, 10mM ß-mercaptoethanol, 1mM MgCl_2,_ 1mM Imidazole, 0.1% NP-40, 4mM CaCl_2_) by slow rotation on a rotator wheel overnight at 4°C. The beads are collected by centrifugation the next day and washed five times with Calmodulin Binding Buffer. Finally the beads are incubated with 200ul of elution buffer for 10 minutes at 4°C by slow rotation on rotator wheel. This process in repeated five times so that the elution volume is 1ml.

### SDS-PAGE, Blue Silver Staining & Western Blotting

For most experiments, an 8% SDS-PAGE gel of 1.5mm thickness was used. About 60-80ug of whole cell extract was mixed with 2x sample loading dye to a final concentration of 1x in a total volume of 20ul and boiled for 5minutes and loaded onto SDS-PAGE gel following which the gel was stained by blue-silver or colloidal Coomassie technique (Dyballa *et. al*, 2009). The bands are visualised under white light using a light-box.

Additionally, gels can also be processed for western blotting in which case transfer of proteins was setup onto a PVDF membrane (PALL Life Sciences). Transfer was carried out at 30V in appropriate transfer buffer for overnight. After transfer, the protein is checked for total protein levels as well as for transfer efficiency by staining with Ponceau S stain at room temperature. The membrane was then blocked with 5% blocking solution in 1x TBST for 1 hour at room temperature, followed by incubation with primary antibody of specified dilution (diluted in 5% BSA) for 1 hour at room temperature. The membrane was washed with 1x TBST four times and then incubated with secondary antibody diluted accordingly in 5% BSA for 1 hour at room temperature. Following this incubation, the membrane was washed again with 1x and the bound antibody was detected with ECL Western Blotting Substrate kit (abcam) and the chemiluminescence captured by exposure to a Carestream XBT X-ray Film. The reagent compositions used for the assay are detailed below.

### Co-Immunoprecipitation

Cells were grown in 50ml YPD until OD_600_ 2.7-3.0 is reached. Cell lysate was prepared as described previously in 1ml of Buffer A. Protein A ^PLUS^ Agarose Beads (BioBharati LifeSciences Pvt. Ltd.) ∼50ul bed volume per reaction were equilibrated twice with ten volumes of Buffer A. The beads for each washing was resuspended in Buffer A and kept on a rotator wheel for 5mins at 4°C. The beads were centrifuged each time to remove residual Buffer A. The extract was then pre-cleared by adding the equilibrated beads and incubation on a rotator wheel at 4°C for 30 minutes followed by centrifugation to get rid of the beads. To the pre-cleared protein extract 2.5ug of anti-TAP Ab (Thermo Fisher Scientific, CAB1001) was added and incubated for 4 hours on rotator wheel at 4°C. To this extract 100ul bed volume of pre-equilibrated Protein A ^PLUS^ Agarose Beads was added and incubated at 4°C on rotator wheel overnight. The beads are washed thrice with ten volumes of Buffer A by rotating on rotator wheel at 4°C for ten minutes each. Finally elution is performed by boiling the beads in 40ul of SDS loading dye for 5 minutes. Samples are now ready to be analysed on SDS-PAGE gel followed by Western blot analysis. Additionally, 50 units of Benzonase Nuclease (Novagen) supplemented with 0.5mM DTT in a total volume of 1ml is added to the pre-cleared protein extract to test for RNA-mediated protein interactions. (Prepare Buffer A without EDTA as EDTA hampers Benzonase Nuclease activity).

### Sample preparation for LC/MS-MS

Cells were harvested at OD_600_ 2.7-3.0 from 2-3 ltrs YPD media at 5000rpm for 5minutes at 4°C and washed once with 10ml of distilled water. Cells were then resuspended in 3ml of Buffer A to which 2.2gms of glass bead was added. Cells were then vortexed for 25 times with 20second pulses with 30 second incubation on ice in between each vortex and centrifugation is performed according to the cell lysate preparation procedure as described above. The lysate is then incubated overnight with 400ul of equilibrated IgG slurry (IgG Sepharose^TM^ 6 Fast Flow from GE Healthcare) at 4°C with slow rotation. The beads are subjected to TEV cleavage as described previously followed by incubation with 600ul pre-equilibrated Calmodulin beads (Calmodulin Sepharose^TM^ 4B from GE Healthcare) in 3ml Calmodulin Binding Buffer by slow rotation on a rotator wheel overnight at 4°C. Final elution is done in 1ml of elution buffer by incubating beads with 200ul of elution buffer for 10 minutes at 4°C by slow rotation on rotator wheel. This eluate is then subjected to LC-MS/MS analysis using Acquity Waters UPLC Systems (Facility run by Sandor Lifesciences Pvt. Ltd.).

### Immunofluorescence

Immunofluorescence experiments were executed as described in Wente *et. al*, 1992. Briefly cells were grown in 10ml culture tubes up to mid-log phase. Cells were now fixed in 4% paraformaldehyde solution for 30 minutes after getting rid of the residual culture media. Cells are next washed twice in 0.1M KPO4 and once in 0.1M KPO4/1.2M Sorbitol and finally resuspended in 1ml 0.1M KPO4/1.2M Sorbitol. Following this cells are spheroplasted for permeabilisation. To about 0.5ml cells 5ul of 10mg/ml zymolase is added and incubated for 10-20mins as per cell density for spheroplasting. The reaction tube is supplemented with 10ul of beta-mercaptoethanol for optimal reaction. After adequate spheroplasting, cells are pelleted at 5000rpm for 1minute and washed twice gently with 0.1M KPO4/1.2M Sorbitol and finally resuspended in 200ul of the same. Cells are now ready to be adhered onto poly-lysine coated cover-slips (regular glass-slips are appropriately poly-lysine coated according to Manufacturer’s protocol). About 100ul of spheroplasted cells are added onto the poly-lysine coated cover-slips and allowed to sit for 15-20 minutes. After aspirating out the supernatant cells are blocked with 400ul of PBS-BSA for 30 minutes followed by the addition of primary antibody (anti-TAP antibody at 1:200). Cells are incubated overnight at 4°C in humid chamber. The cells were then washed five times with PBS-BSA before incubation with secondary antibody (Goat anti-rabbit secondary antibody, Alexa Fluor® 488 at 1:100) at room-temperature in the dark for 1hour. The cells were then washed extensively with PBS-BSA and finally mounted onto a clean glass slide using 10mg/ml p-phenylenediamine (Sigma) in 90% glycerol which is already supplemented with DAPI (20ug/ml). The cells were observed in Confocal Laser Scanning Microscope (Olympus, Model-IX81) and the localization of eIF4G analyzed. The reagents used for this assay are detailed below.

## Results

### The N-terminally located RRM1 domain of Tif4631p play an indispensable role in the nuclear mRNA surveillance of aberrantly long 3′-extended and export inefficient messages

Tif4631p (also known as the eukaryotic translation initiation factor gamma or eIF4G) is an ubiquitous protein that plays a crucial functional role in the regulation of cap-dependent translation in eukaryotes (Pyronnet et al. 1999; Hershey et al. 2012). Although, this protein was initially identified as a cytoplasmic factor with a vital cytoplasmic function, several recent findings demonstrated various nuclear functions such as splicing (Kafasla et al. 2009), nuclear mRNA degradation and surveillance (Das et al. 2014), and possibly a nuclear translation (Iborra et al. 2004). Considering its role in diverse nuclear and cytosolic RNA metabolic events, Tif4631p was previously hypothesized to serve as an adapter of various protein complexes (Das and Das 2016) involved in splicing of pre-mRNAs (Kafasla et al. 2009), nuclear mRNA surveillance(Das et al. 2014), and initiation of translation (Goyer et al. 1993; Tarun Jr. and Sachs 1996, 1997; Wells et al. 1998). Biochemical evidences suggest that yeast Tif4631p harbours following functional domains from the N-terminus to C-terminus (Fig. 1) that include RRM1 (RNA recognition motif) followed by Pab1p-binding, Cbc1p-binding (overlapping with RRM2), MIF4G (N-terminal end of this domain overlaps with the C-terminal end of Cbc1p-binding domain) and RRM3 (Tarun and Sachs 1996, 1997; Altmann *et al*. 1997; Dominguez *et al*. 1999, 2001; Asano *et al*. 2001) (Fig. 1). Notably, MIF4G (*M*iddle Domain of e*IF4G*), the most prominent domain comprising of 240 amino acid residues is located toward the C-terminal region (overlapping with eIF4A-interacting domain). Previous bioinformatics analysis revealed that this domain presumably adopt an *-*helical structure and may participate in both protein–protein and protein–RNA interactions (Aravind and Koonin 2000; Ponting 2000). The MIF4G domain was implicated as a multipurpose adaptor, which potentially connects the two other elements of the eIF4F complex, eIF4A and eIF4E (Lamphear *et al*. 1995; Imataka and Sonenberg 1997; Craig *et al*. 1998), and to eIF3, the 40S ribosomal subunit and mRNA (Pestova, Shatsky and Hellen 1996). Three RRMs are located at the N-terminal, middle and C-terminal regions respectively. The latter two are rich in arginine and serine (Lomakin, Hellen and Pestova 2000; Berset *et al*. 2003; Prevot, Darlix and Ohlmann 2003; Shatsky *et al*. 2014). We addressed the functional contribution of each of these domains in the surveillance of select aberrant and special mRNAs by evaluating the ability of various deletion clones of Tif4631p each lacking a specific domain (Fig.1A and Table 1) to target and degrade the select aberrant and special nuclear mRNAs. In addition, we also assessed the domain contribution by carrying out two different genetic assays – one consisting of the suppression of genetically constructed *LCY* allele and suppression of *lys2-187* mutation in an SC-Lys medium (see below).

**Figure 1.**
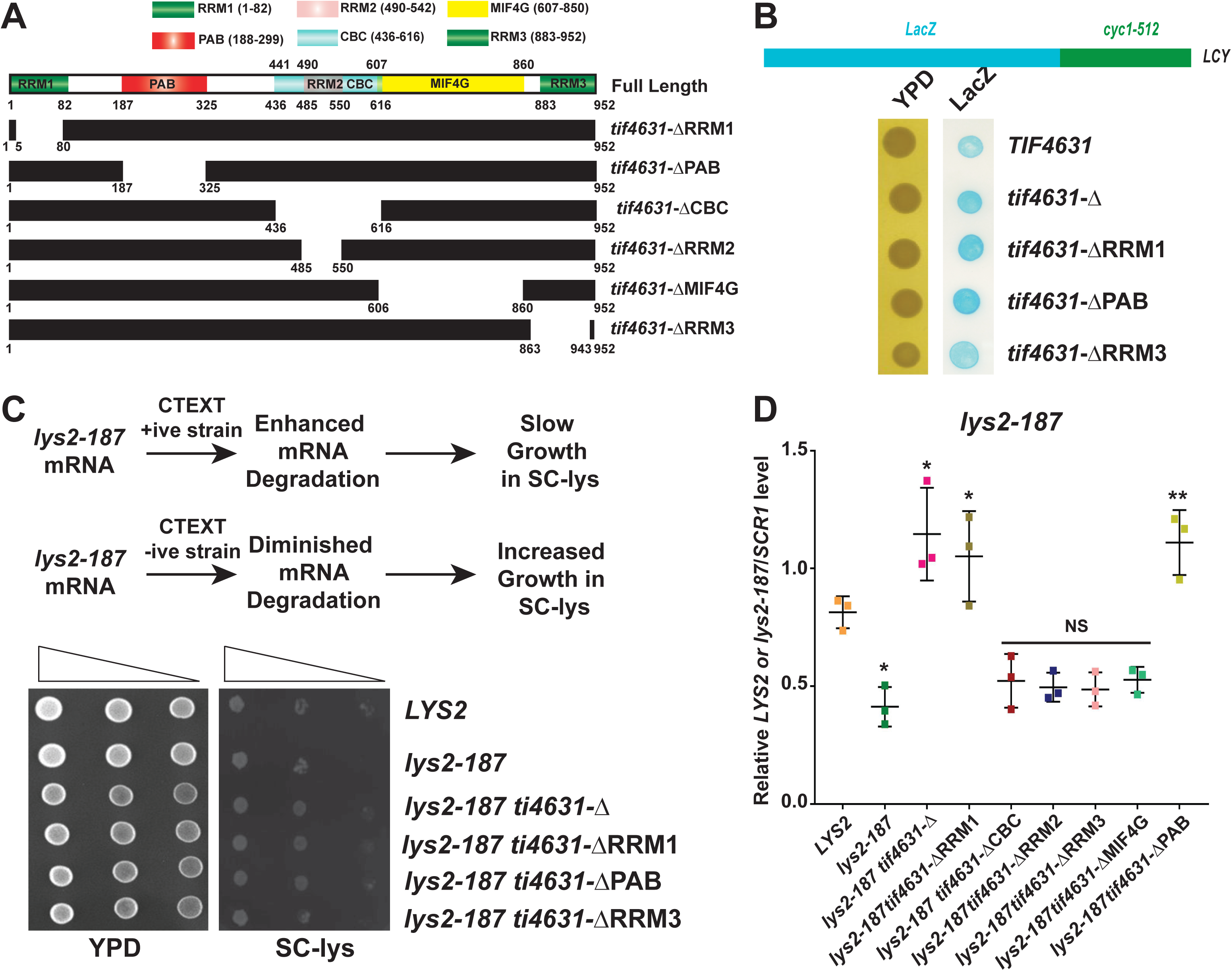
The RRM1 domain (1-82 aa) of Tif4631p contributes actively to the degradation of DRN/Exosome substrate mRNAs. (A) A schematic representation of Tif4631p domain deletions along with their demarcated amino acid boundaries adopted for this study. (B) Schematic representation of *LCY* allele. Growth phenotype of WT, *tif4631-*Δ*, tif4631-* Δ*RRM1, tif4631-*Δ*PAB, tif4631-*Δ*RRM3*, was tested on lactose media supplemented with X-gal (20 mg/ml). Equal number of cells from indicated strains was spotted on YPD and lactose media supplemented with X-gal (20 mg/ml) simultaneously. Cells are then grown at 30°C for sufficient period and monitored regularly. (C) Suppression of growth of *lys2-187* allele by various domain deletion mutants of Tif4631p. Schematic representation of the mechanistic rationale behind the plate assay experiment. Growth of *lys2-187* yeast strain is significantly suppressed by *tif4631-*Δ (yBD 214) on *LYS* dropout medium. yBD 191 carrying a perfectly healthy allele of the *LYS2* gene grows successfully on the same dropout media and serves as the positive control for this assay, while yBD 20 carrying mutant *lys2-187* allele fails to grow on the dropout media and serves as the negative control for this experiment. The RRM1 domain deletion (yBD 387) and PAB domain deletion (yBD 389) produces similar suppression of the mutant *lys2-187,* with cells growing successfully on *LYS* dropout medium. The RRM3 domain deletion (yBD 420) fails to grow on the dropout media thus confirming no contribution of this domain in *lys2-187* turnover in the nucleus. Equal number of cells of *LYS2^+^*(yBD 191), *lys2-187* (yBD 20), *tif4631-*Δ (yBD 214), *tif4631-*ΔRRM1 (yBD 387) and *tif4631-*ΔPAB domain deletion (yBD 389) were spotted with ten-fold dilution on *LYS* dropout media. Cells are then grown at 30°C for sufficient period and monitored regularly for growth. (D) Relative steady state levels of export inefficient *lys2-187* mRNA at 30°C in *LYS2*, *lys2-187 lys2-187 tif4631-*Δ and yeast strain carrying deletions in RRM1, CBC1, RRM2, RRM3, MIF4G and PAB domains. The *upf1-*Δ was used as negative control. Normalized value of specific mRNA from wild-type samples was set to 1. For each strain three independent samples of random primed cDNA (biological replicates, N=3) were prepared from strains grown at 30°C and subjected to Real-time PCR analysis using primer sets specific for each mRNA. Transcript copy numbers/2 ng cDNA of each strain were normalized to *SCR1* RNA levels in respective strains and are shown as means ± SE. The statistical significance of difference as reflected in the ranges of p-values estimated from Student’s two-tailed t-tests for a given pair of test strains for each message are presented with following symbols, * <0.05, **<0.005 and ***<0.001, NS, not significant. (D)

We initiated our studies by assessing if the decay of 3′-extended mRNAs undergo decay in the yeast strains carrying various clones having a specific deletion in each domain (Fig 1A, Table 1). To carry out this experiment we employed a genetically engineered *LCY* (*<u>L</u>acZ-<u>CY</u>C1*) allele. This allele was constructed by fusing the ORF of the *LacZ* to the 3′-UTR of *cyc1-512* gene (Zaret and Sherman 1982, 1984; Das et al. 2000)(Fig. 1B). The *cyc1-512* mutation consisting of a 38-bp deletion at eight nucleotides (nt) upstream from the normal poly(A) site led to the elimination of the signals required to form transcription termination and mature 3′-end of yeast mRNAs (Zaret and Sherman 1982, 1984). Consequently, abnormal transcription termination and transcription read-through the *cyc1-512* allele resulted in the formation of eight aberrantly long *cyc1-512* mRNAs discrete 3′-termini ranging from the wild-type poly(A) end-points to the site with an endpoint that is more than 3500 nt long (Zaret and Sherman 1982, 1984; Das et al. 2000). These aberrant mRNAs are subjected to the rapid decay by the nuclear exosome and DRN (Das et al. 2000). Since, it is difficult to determine the stability of the aberrantly long *cyc1-512* mRNAs, we employed a novel genetic assay using the *LCY* allele to qualitatively assess the extent of decay of aberrantly long 3′-extended *LCY* mRNA. Notably, owing the presence of 3′-UTR of *cyc1-512* gene, the transcription in *LCY* allele continued beyond the 3′-end of the *LacZ* gene and found to produce aberrantly long 3′-extended *LCY* mRNAs. These aberrantly long *LCY* mRNAs were subjected to rapid degradation by DRN and the nuclear exosome (Das et al. 2000) thereby producing reduced amount of β-galactosidase. An impairment in the DRN or exosome machinery consequently stabilized these aberrant mRNAs due to diminished decay that led to the increased production of β-galactosidase. Thus a yeast strain expressing an *LCY* allele and the full length Tif4631p is expected to yield very faint blue colonies when grown in *IPTG* containing medium (Fig. 1B). In contrast, a strain carrying a deletion in *TIF4631* gene (*tif4631-*Δ) expressing the same *LCY* allele is predicted to yield a darker blue colony due to diminished nuclear decay of *LCY* mRNAs due to impaired nuclear mRNA decay machinery when grown in the same IPTG containing medium (Fig. 1B). Subsequent evaluation whether any of the domain deleted mutants of Tif4631p impairs the rapid degradation of the *LCY* mRNAs revealed that both of the *tif4631*-ΔRRM1 and *tif4631*-ΔPAB mutants yielded darker blue colony like the *tif4631-*Δ mutant (Fig.1B), thereby indicating that the RRM1 and PAB domains may play a crucial role in the nuclear decay of aberrant 3′-extended messages.

Next, we went a step forward to evaluate if these same sets of deletion in Tif4631p could result in a similar suppression of rapid decay of mRNA export-defective messages. Different categories of export-defective mRNAs are generated during the final phase of nuclear mRNP biogenesis in baker’s yeast that were previously categorized in three different categories (Maity et al. 2016). These include (i) those that are caused by a *cis*-acting mutation in their transcript body as exemplified by export-defective *lys2-187* mRNA (Das et al. 2006) (ii) global poly(A)^+^ mRNAs arrested in the nucleus of the temperature-sensitive export-mutant *nup116*-Δ strain at conditional temperature of 37°C due to complete block of the nuclear export (Das et al. 2003) and (iii) naturally occurring nucleus-retained export-inefficient non-aberrant special mRNAs (Kuai et al. 2005). All three categories of export-defective messages undergo rapid degradation in the exosome and DRN-dependent and TRAMP-independent manner (Maity et al. 2016). Our quest concerning the effect of the deletion of specific domain(s) of the Tif4631p on the rapid degradation of export-defective mRNAs began with the evaluation of the suppression pattern of *lys2-187* allele. The *lys2-187* allele contains two base substitutions (one of them is located at 1052 and the other is located at 1054 nt of *LYS2* open reading frame (Das et al. 2006). This mutation causes a structural alteration of the corresponding *lys2-187* mRNA leading to its nuclear retention and a rapid degradation by the nuclear exosome and DRN and a very slow growth of a *lys2-187* strain (relative to an isogenic *LYS2* strain) on SC-lys medium (Das et al. 2006; Maity et al. 2016) (Fig. 1C). A mutation in any component of either of the core exosome or DRN led to the stabilization of this message resulting in the relatively rapid growth on the SC-lys medium (Das et al. 2006; Maity et al. 2016) (Fig. 1A). A close examination of the ability of the growth of the *tif4631-*Δ yeast strain and the strains carrying the specific deletion of the various functional domains in Tif4631p (Fig. 1A, Table 1) showed that the *lys2-187* allele was suppressed by the yeast strains expressing *tif4631-*Δ, *tif4631*-ΔRRM1 and *tif4631*-ΔPAB deletion clones (Fig. 1C) suggesting that perhaps the rapid decay of *lys2-187* mRNA may be stabilized by these mutations. To further bolster this finding, we measured the steady-state levels of the *lys2-187* message as an indicators of its decay rates in *LYS2*, *lys2-187*, *lys2-187 tif4631-*Δ and strains harboring various domain deletion in Tif4631p proteins. As shown in Fig. 1D, the steady-state levels of the *lys2-187* mRNA is significantly low in *lys 2-187* strain, relative to *LYS2* strain (Fig.1D). Under the same condition, the levels of *lys2-187* mRNA was enhanced in strains carrying *tif4631*-ΔRRM1 and *tif4631*-ΔPAB deletions. This data therefore, clearly corroborated the data presented in Fig 1B and 1C). The other deletion clones such as *tif4631*-ΔRRM3/2 and *tif4631*-ΔCBC deletion clones were unable to cause any increase in the steady-state levels of *lys2-187* mRNA (Fig. 1D). This observation strongly supports the notion that the RRM1 and PAB domains of Tif4631p may be major contributing domain in the nuclear surveillance/degradation of export defective *lys2-187* mRNA.

Finally, we addressed whether the decay/steady-state levels of other categories of export defective messages as exemplified by special mRNAs as well as the global poly(A) RNA trapped in the nucleus of *nup116*-Δ strain are also similarly affected. The steady-state levels of two special mRNAs, *IMP3* and *NCW2* displayed an enhancement of about 2.5 to 4 folds in yeast strains carrying a *tif4631*-Δ, *tif4631*-ΔRRM1 and *tif4631*-ΔPAB deletion strains relative to corresponding yeast strain carrying a *TIF4631* allele. Curiously, the extent of enhancement of the steady-state level of *IMP3* message in *tif4631*-ΔRRM1 was found to be much lower relative that observed for *tif4631*-ΔPAB strain (Fig. 2A), the reason of which was not clear to us. Moreover, the levels of these special mRNAs did not change appreciably in the yeast strains expressing the other deletions of Tif4631p (Fig. 2A). Under the same conditions the steady-state levels of two non-special typical mRNAs, *ACT1* and *CYC1*, in contrast did not alter in any of the indicated yeast strains (Supplementary Fig. S1 B-C). Furthermore, we observed that both the levels of two representative poly(A)^+^ RNAs, *ACT1* and *CYC1* were significantly lower in *nup116*-Δ *TIF4631* strain at the conditional temperature of 37°C relative to that observed in the corresponding isogenic *NUP116 TIF4631* strain (Fig. 2B). This finding is supportive of the fact that both of these mRNAs undergo a rapid and active degradation under the condition of complete retention in the nucleus that occurs due to complete block of their nuclear export at 37°C, which is consistent with the previously published observation (Das et al. 2003). Notably, their nuclear decay is dependent on the exosome and DRN since depletion of the components of any of these machinery displayed a significant stabilization of both of them as depicted by the depletion of DRN component Tif4631p (*tif4631-*Δ strain) (Fig 2B). Interestingly, their steady state levels were found be enhanced by about 3 to 4 folds in *nup116-*Δ yeast strains additionally carrying a *tif4631*-ΔRRM1 and *tif4631*-ΔPAB deletion relative to *nup116-*Δ *TIF4631* strain (Fig. 2B). Importantly, the levels of these two mRNAs estimated from the indicated yeast strains grown at the permissive condition of 25°C did not exhibit any significant alteration in any of these strains (Supplementary Fig S2 B-C) since being a temperature-sensitive mutation, *nup116*-Δ does not display any export defect at 25°C and global mRNAs undergo efficient export thereby did not cause any nuclear arrest of any of the poly(A)^+^ RNA (Das et al. 2003). This finding, thus, suggests that the N-terminally located RRM1 and PAB1 domains play crucial roles in the degradation of both representative special mRNAs and nucleus-trapped poly(A) RNAs.

**Figure 2.**
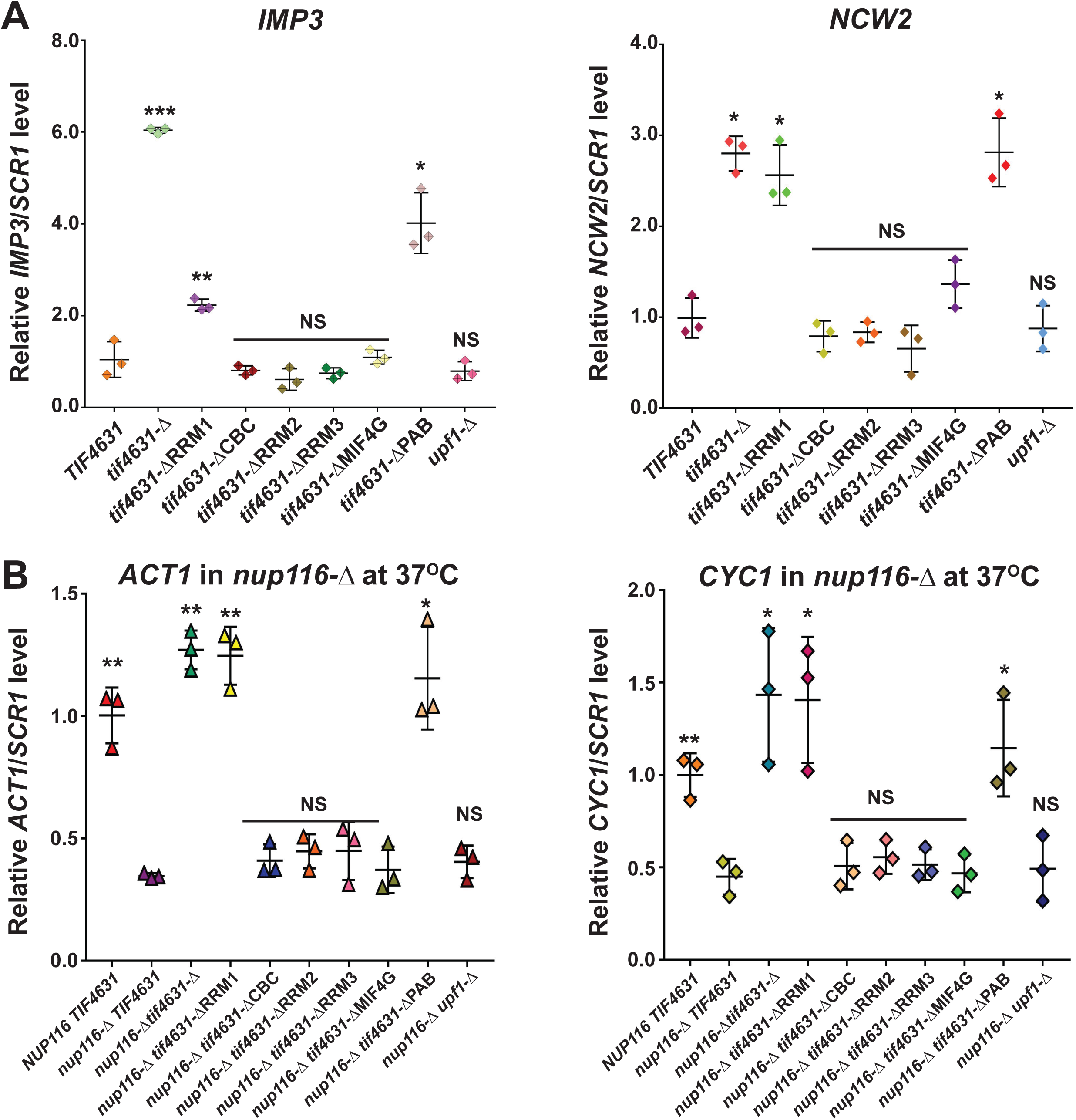
The RRM1 domain (1-82 aa) of Tif4631p contributes actively to the degradation of export defective mRNAs. Relative steady state levels of export inefficient special mRNAs *IMP3* & *NCW2* at 30°C (A) and *ACT1* & *CYC1* mRNAs trapped in the nucleus of *nup116*-Δ yeast strain background (B) in *TIF4631*, *tif4631-*Δ and yeast strain carrying deletions in RRM1, CBC1, RRM2, RRM3, MIF4G and PAB domains. The *upf1-*Δ is used as negative control. Normalized value of specific mRNA from wild-type samples was set to 1. For each strain three independent samples of random primed cDNA (biological replicates, N=3) were prepared from strains grown at 30°C and subjected to Real-time PCR analysis using primer sets specific for each mRNA. Transcript copy numbers/2 ng cDNA of each strain were normalized to *SCR1* RNA levels in respective strains and are shown as means ± SE. The statistical significance of difference as reflected in the ranges of p-values estimated from Student’s two-tailed t-tests for a given pair of test strains for each message are presented with following symbols, * <0.05, **<0.005 and ***<0.001, NS, not significant.

Thus, the data presented in Figs. 1 and 2, is extremely consistent in that the deletion of both the N-terminal RRM1 and PAB domains caused an enhancement of the steady-state levels of both the aberrantly long 3′-extended and export-inefficient messages (Figs. 1 and 2). In order to rule out the possibility, that the enhancement of stead-state levels of these select mRNAs in strains carrying *tif4631-*ΔRRM1 and *tif4631*-ΔPAB alleles is due either to the decreased expression or to the altered nuclear localization/import of these deletion versions of Tif4631p, we also evaluated the levels of deletion versions of Tif4631p (Tif4641p-ΔRRM1 and Tif4641p-ΔPAB) by western blot analysis from the extract prepared from these strains and compared with that of the full length Tif4631p. As presented in Supplementary Fig. S2 D, although the level of and *tif4631*-ΔRRM1 protein was found to be comparable to that of the full length protein, the expression level of *tif4631*-ΔPAB protein could not be detected (Fig. S2D), the reason of which was not clear to us. Therefore, we are unable to draw any conclusion concerning the effect of the *tif4631*-ΔPAB protein on the stability and decay of any of the aberrant mRNAs. Furthermore, we consequently examined the *in situ* cellular distribution of full length Tif4631p and *tif4631*-ΔRRM1 deletion protein using indirect immunofluorescence. As shown in supplementary figure S2E, the cellular localization and distribution pattern of both of them were essentially similar showing they are distributed both in the nucleus and cytoplasm that is consistent with the previously published data (Kafasla et al. 2009). These results, thus, together suggestive of the conclusion that the N-terminal RRM1 domain of the Tif4631p plays a seminal functional role in the nuclear degradation/surveillance of the aberrantly long 3′-extended and export-defective messages in baker’s yeast. However, to get into the bottom of this finding we address the exact molecular role of this protein as well as this domain (see next section).

### Tandem Affinity Purification using Tif4631p as bait leads to the co-purification of previously identified components of DRN, the nuclear exosome and TRAMP

Having assigned a functional role of the N-terminal RRM1 domain of Tif4631p, we stepped forward to evaluate if Tif4631p (and this domain) plays a crucial functional role in the complex formation of DRN proteome and/or in mediating the interaction of DRN component with the core nuclear exosome. We initiated this effort by accomplishing a comprehensive definition of DRN and by addressing if the genetically identified DRN components display any physical association/ interactions among themselves. We purified the C-terminally TAP-tagged Tif4631p from a yeast strain using a tandem affinity purification procedure and identified the interacting proteins by mass-spectrometry (Puig et al. 2001) (Fig. 3B-D). We chose Tif4631p as the bait in this co-purification experiment, since it was previously hypothesized to play the role of an integrator of various protein complexes involved in several aspects of RNA metabolism (Das and Das 2016). We consistently and reproducibly identified Cbc1p, Dbp2, Upf3, Nrd1p (NNS component), Mtr4p (component of TRAMP) and Rrp4p (component of the nuclear exosome) co-purified with Tif4631p. Among them, Cbc1p (Das et al. 2000, 2003, 2006), Upf3p and Tif4631p (Das et al. 2014) were previously identified and demonstrated to take part in the degradation of 3′-extended aberrant and export defective messages (Maity et al. 2016). Strikingly, Cbc1p and Upf3p appeared to be two most abundantly co-purifying proteins (Fig. 3D), which is consistent with the conclusion stemmed from the previous genetic data that together with Tif4631p, Cbc1p and Upf3p collectively function in the nuclear mRNA surveillance possibly by forming a stable protein complex. Interestingly, a new Tif4631p-inetracting protein, an ATP-dependent DEAD box RNA helicase, Dbp2p (Cloutier et al. 2012; Ma et al. 2013; Lai et al. 2019) also co-purified with Tif4631p as a strong interacting partner in a reproducible manner. Subsequent genetic experiments suggest that DRN-specific select export-defective mRNAs undergo significant stabilization in a yeast carrying a deletion in *DBP2* gene (see next section, Fig. 6). This data strongly suggests that Dbp2p is possibly a newly identified component of DRN system. In a good agreement with our finding, Dbp2p was previously demonstrated to participate in the nuclear mRNA surveillance (Cloutier et al. 2012) and cytoplasmic nonsense-mediated mRNA degradation and rRNA biogenesis (Bond et al. 2001). Remarkably, other than these four proteins, NNS complex protein, Nrd1p; core-nuclear exosome component, Rrp4p; and TRAMP component Mtr4p also co-purified in our TAP affinity purification procedure followed by LC-MS/MS analysis (Fig. 3A and D). This finding is perfectly consistent with our expectation and previous model that together with the nuclear exosome and TRAMP, DRN target the aberrantly long 3′-extended messages and along with the nuclear exosome, degrade the export inefficient messages (Das et al. 2000, 2003, 2014; Maity et al. 2016). Consequently, known DRN components displayed genetic interactions with the components of the nuclear exosome and TRAMP (Das et al. 2000, 2003, 2014; Maity et al. 2016). Furthermore, the co-purification procedure also yielded numerous splicing factors as well as factors involved in translation initiation and cytoplasmic decay factors (Fig. 3D). This observation also makes a perfect sense since Tif4631p was previously reported to participate in splicing of a small subset of mRNAs (Kafasla et al. 2009) and are very well known to regulate the initiation event of the cap-dependent translation (Tarun Jr. and Sachs 1997; Tarun Jr. et al. 1997; Wells et al. 1998; Hilbert et al. 2011). Thus, the co-purification of the previously known interacting partners of Tif4631p during our TAP-purification procedure strongly validate our LC-MS/MS data and thereby imparting a significance and relevance to the newly identified interaction of Tif4631p with the Cbc1p, Upf3p and Dbp2.

**Figure 3.**
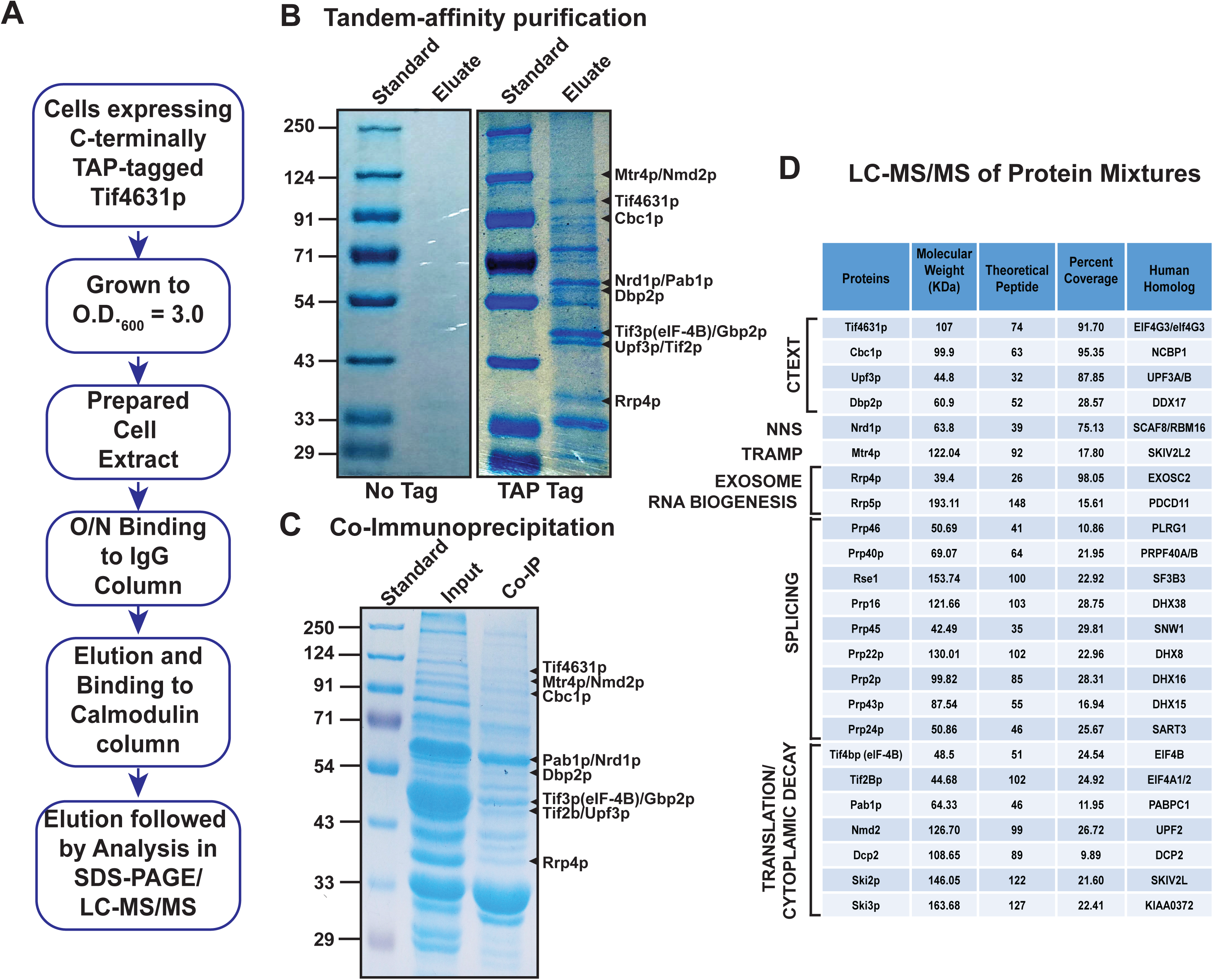
Components of the DRN co-purify with Tif4631p. (A) Schema depicting the rationale of purification of C-terminally TAP-tagged Tif4631p in IgG and calmodulin columns in tandem. The eluate is analyzed simultaneously by SDS-PAGE and LC-MS/MS. (B) Representative pulled-down proteins using the experimental rationale outlined in (A), with untagged and Tif4631p-TAP tagged strains. The pulled down proteins were compared side by side on SDS-PAGE followed by blue-silver staining. A pre-stained standard was electrophoresed alongside for size comparison of proteins present in the eluate. Proteins identified mass spectroscopy are indicated by arrows (C) Representative pull-down proteins from immunoprecipitation using TAP-tagged Tif4631p as bait were analyzed side by side on SDS-PAGE followed by blue-silver staining. Proteins identified mass spectroscopy are indicated by arrows (D) Table depicting the results of liquid-chromatography tandem mass-spectroscopy (LC-MS/MS) analysis of the eluate from the IgG and calmodulin columns used in tandem using Tif4631p-TAP as bait. List of proteins that co-purified with Tif4631p-TAP showing the theoretical peptide and percent coverage values associated with each one and their corresponding human homologs.

To further bolster this finding, we also subjected the protein extract prepared from the yeast expressing Tif4631p-TAP to the immunoprecipitation using an anti-TAP antibody (Thermo Scientific) followed by the analysis of the co-immunoprecipitate either by gel or by LC-MS/MS, which displayed a very similar pattern of the protein in the immunoprecipitate (Fig. 3C). Even in this approach, we observed Cbc1p, Upf3p, and Dbp2p to reproducibly co-purify when the protein extract prepared from the yeast strain expressing Tif4631p-TAP was precipitated using the TAP antibody. Thus, both independent purification schemes yielded a very similar and consistent profiles of the interacting partners of Tif4631p thereby very strongly indicating that Tif4631p, Cbc1p, Upf3p and Dbp2p form a stable complex.

### Systematic proteomic analysis establishes a physical association among components of DRN, and the core nuclear exosome

Having identified Tif4631p-inetracting proteins by LC-MS/MS, we then ask whether a similar co-purification profile would result if each of the Tif4631p-interacting proteins is individually used as bait in an immunoprecipitation experiment linked to their detection. Consequently, we first validated the LC-MS/MS data presented in Fig. 3 by subjecting the protein extract prepared from yeast strain expressing Tif4631p-TAP to immunoprecipitation using TAP antibody. The resulting immunoprecipitate was sequentially analyzed by western blotting using antibodies against various proteins as shown in the scheme presented in Fig. 4A. As shown in Fig, 4B and C, when Tif4631p was used as bait to immunopurify its interacting partners, all the previously identified proteins by LC-MS/MS, Cbc1p, Upf3p, Dbp2, as well as the NNS component Nrd1p and exosome component Rrp4p and Rrp6p could be detected (Fig. 4B-C) thereby corroborating our original LC-MS/MS result. We also tested the RNA dependence of these interactions by treating the cell extracts with benzonase nuclease overnight before subjecting it to the immunoprecipitation (Fig. 4A and C). As shown in Fig. 4C, interactions of Tif4631p with Nrd1p and Rrp6p appeared to greatly diminish upon benzonase treatment, thereby indicating that possibly the Tif4631p-Nrd1p and Tif4631p-Rrp4p interaction require the presence of RNA. This finding is consistent with the idea that Nrd1p and Rrp4p belong to NNS and core-exosome complex respectively, which are not a part of the integral component of Tif4631p-dependent complex. Taken together, we favor the view that together, Tif4631p, Cbc1p, Upf3p and Dbp2p form an integral complex.

**Figure 4.**
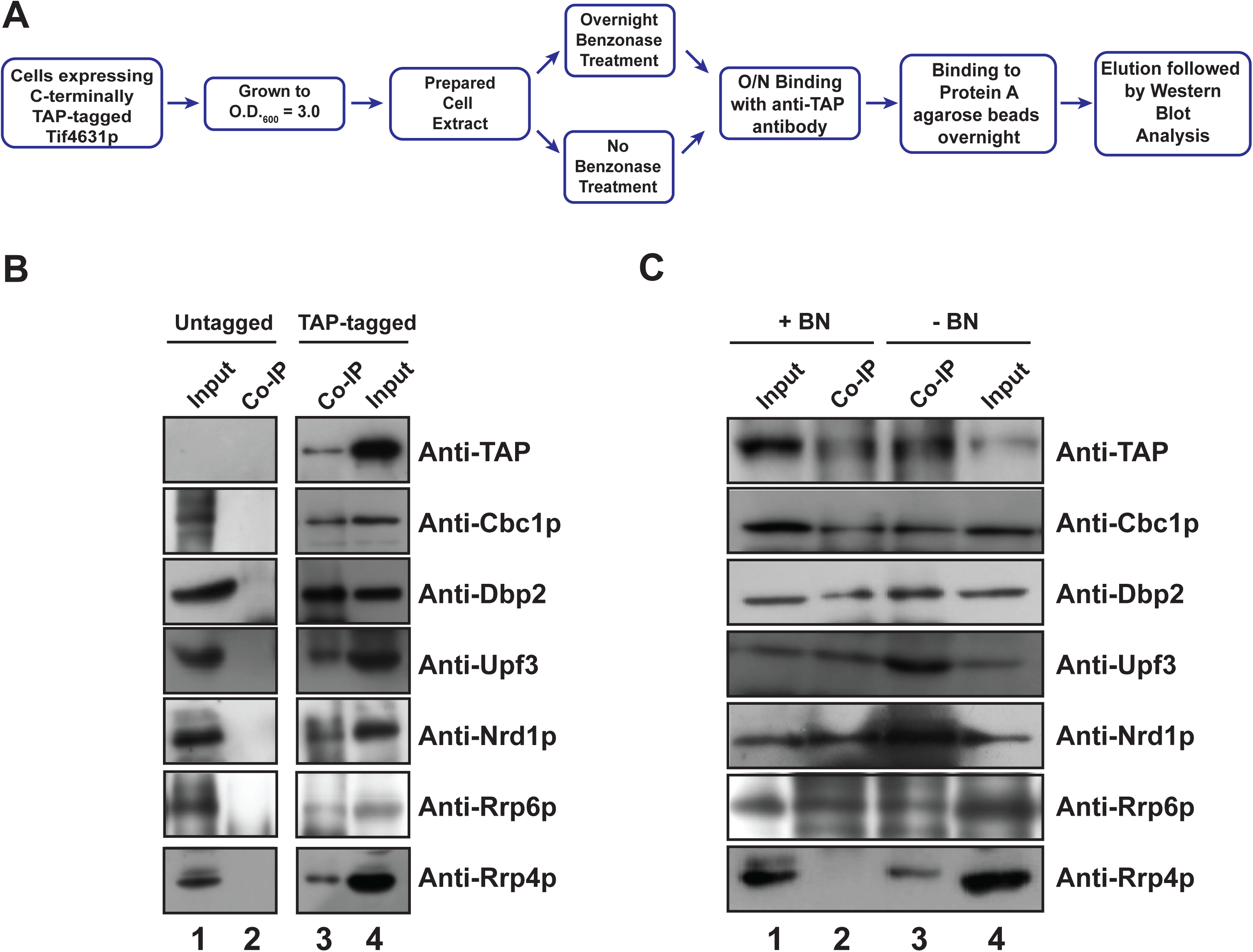
The CTEXT complex consists of Tif4631p, Cbc1p, Upf3p and Dbp2p as core components which interact with components of the core nuclear exosome and Nrd1p. (A) Schematic outline of the co-immunoprecipitation experiment using Tif4631p-TAP as bait. (B) Result from a representative co-immunoprecipitation experiment using an untagged and TAP-tagged Tif4631p as bait. Protein extracts from untagged Tif4631p and genomically TAP-tagged Tif4631p are processed according to the protocol in Fig. 4A, electrophoresed on denaturing SDS gel followed by the detection of the specific CTEXT components using western blotting by using specific antibodies. The input (Lanes 1 and 4) in case of each sample is 100 μg of whole-cell protein extract from the respective strains, while the Co-IP (Lanes 2 and 3) for each lane represents 20 μl of the eluate in SDS-gel loading buffer from respective strains. The untagged strain acts a negative control while the anti-TAP lane serves as the positive control for this set of experiments. (C) Result from a representative co-immunoprecipitation experiment using TAP-tagged Tif4631p as bait followed by treating the IP either with benzonase nuclease (+BN) or without any treatment (-BN). Detection of specific CTEXT components were then done by western blotting as described above.

To further corroborate this finding, we extend our investigation to evaluate if same sets of interacting partners co-purify in a reproducible and consistent manner when other proteins than Tif4631p is used as a bait in separate and independent experiments. As shown in Fig. 5A-B, when either of the Cbc1p-TAP or Dbp2p-FLAG was used as bait in similar independent immune-precipitation procedure, same sets of the proteins were identified by western blotting analysis (Fig. 5A-B). Testing if these interactions require the presence of RNA, we also treated the protein extracts prepared from yeast strains expressing either of the Cbc1p-TAP or Dbp2p-FLAG overnight with benzonase before subjecting them to immunoprecipitation with either TAP or FLAG antibody. Surprisingly, we did not note any RNA-dependence of any of these interactions as the signals of none of these interacting protein did undergo any diminution (Fig. 5A-B). This finding is rather surprising as we expected from the results obtained in experiment involving Tif4631p-TAP as bait (Fig. 4) that the interactions of Nrd1p, Rrp6p, and Rrp4p, with Cbc1p and Dbp2p would be RNA dependent, since these proteins belong to different protein complexes than Tif4631p-driven complex. However, the RNA-dependent interactions of Nrd1p and Rrp6p with Tif4631p may not be a prerogative that the NNS and core exosome would interact with the other components of Tif4631p-driven complex in an RNA-dependent manner. Nevertheless, all these proteomic data, collectively suggest that Tif4631p forms a stable integral protein complex that consists at least of Cbc1p, Upf3p and Dbp2p. We term this complex **CTEXT** (**<u>C</u>**bc1p-**<u>T</u>**if4631p-dependent **<u>EX</u>**osomal **<u>T</u>**argeting) that serves to degrade the 3′-extended aberrant and export-inefficient transcripts in the nucleus by acting as a co-factor of the nuclear exosome (Maity et al. 2016). Consistent with our findings, all the members of CTEXT either known to bind RNA or demonstrated to have at least one RNA recognition motif (RRM), which enable this complex to mediate the degradation of the respective RNA substrates (see below). Henceforth, we redesignate this co-factor complex as CTEXT, which will replace our previously termed name ‘DRN’.

**Figure 5.**
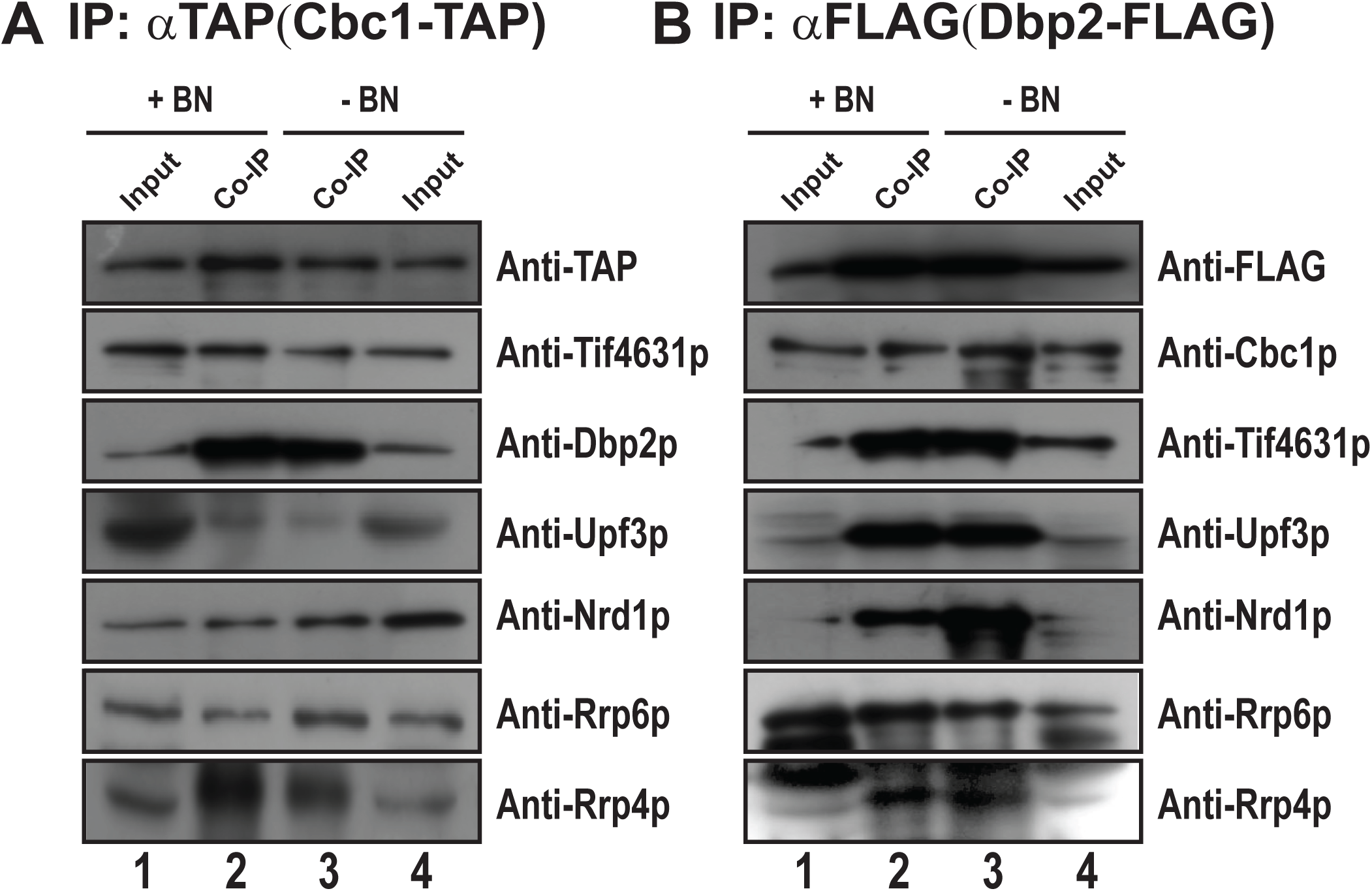
Systematic proteomic analysis establishes a physical association between components of the CTEXT, the core nuclear exosome and Nrd1p. Results of representative co-immunoprecipitation experiment performed with C-terminally TAP-tagged Cbc1p (A) or C-terminally FLAG-tagged Dbp2p (B) as baits followed by treating the IPs from each one either with benzonase nuclease (+BN) or without any treatment (-BN). Detection of specific CTEXT components were then done by western blotting as described above. The anti-TAP or anti-FLAG panels served as the positive control for these experiments. The input in case of each sample is 100 μg of whole-cell protein extract, while the Co-IP for each lane represents 20 μl of the eluate in SDS-gel loading buffer.

### Dbp2p defines a novel component of the CTEXT

Dbp2p, the newly identified member of the CTEXT complex encodes an ATP dependent RNA helicase (Barta and Iggo 1995; Bond et al. 2001) that was previously demonstrated to participate in the remodeling of RNA-protein complex to promote efficient transcription termination (Cloutier et al. 2012), nonsense mediated mRNA decay and rRNA processing (Bond et al. 2001) and plays a key role in the formation of export-competent nuclear mRNPs by enabling the assembly of Yra1p, Nab2p and Mex67p with the maturing mRNAs (Ma et al. 2013). Strikingly, this protein was reported to alter the RNA secondary structure by promoting the unwinding of RNA duplex *in vivo* to promote transcription termination and thereby modulating the proper gene expression (Xing et al. 2017; Lai et al. 2019). However, no previous report indicated that Dbp2p might also be involved in the nuclear mRNA surveillance. To evaluate if this RNA helicase plays an active role in the nuclear decay of the select export-defective mRNAs, we addressed if the deletion of this protein has any effect on the steady-state levels of these substrate mRNAs. As shown in Fig. 6A-C, the steady-state levels of two export-defective special mRNAs, *IMP3* and *NCW2* (Kuai et al. 2005; Maity et al. 2016) and aberrant export-defective *lys2-187* mRNA (Das et al. 2006) displayed a significant enhancement in their steady-state in yeast strain harboring *dbp2-*Δ allele that is comparable with enhancement level shown by the deletion of the known CTEXT member *TIF4631* (*tif4631-*Δ). Notably, no significant increase in the steady-state levels of *ACT1* and *CYC1* mRNAs in the *dbp2-*Δ and *tif4631-*Δ yeast strain was observed. This data thus clearly argue that Dbp2p participates in the nuclear degradation of these mRNAs by specifically targeting them and therefore constitute an essential component of CTEXT complex.

**Figure 6.**
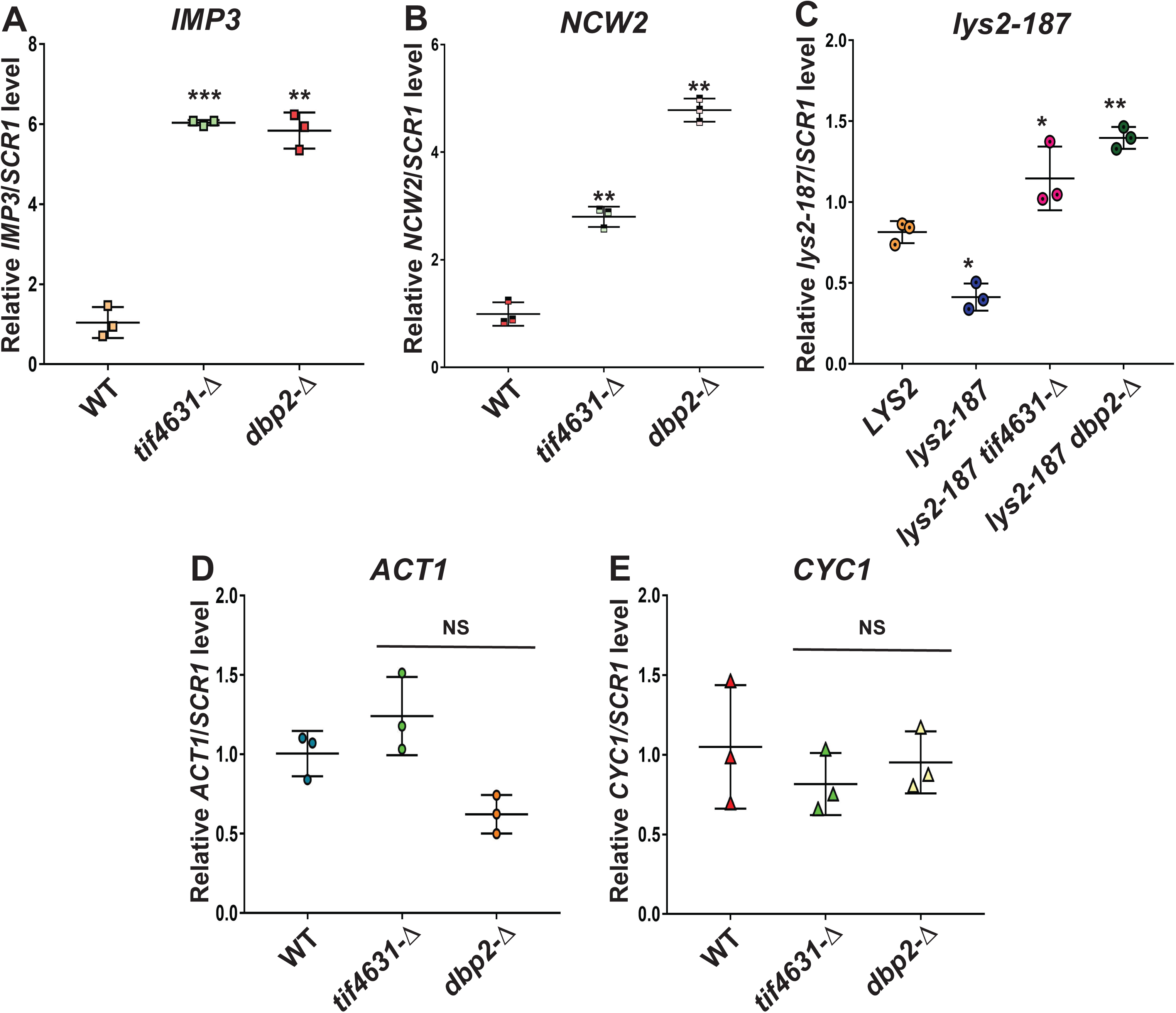
*DBP2,* an ATP-dependent DEAD-box RNA helicase is an important component of the CTEXT proteome. Relative steady state levels of two representative export inefficient special mRNAs (A) *IMP3*, and (B) *NCW2* mRNAs at 30°C in yeast strains carrying *TIF4631*, *tif4631-*Δ & *dbp2-*Δ alleles. Normalized value of specific mRNA from wild-type samples was set to 1. (C) Relative steady state levels of normal *LYS2* and export-defective *lys2-187* mRNA at 30°C in a strain carrying *LYS+TIF4631*, *lys2-187TIF4631*, *lys2-187tif4631-*Δ & *lys2-187dbp2-*Δ. Normalized value of *lys2-187* transcript from the *LYS2+* sample was set to 1. D-E Relative abundance of two representative typical *CYC1* and *ACT1* mRNAs at 30°C in yeast strains carrying *TIF4631*, *tif4631-*Δ & *dbp2-*Δ alleles. Normalized value of specific mRNA from wild-type samples was set to 1. For data presented in Panel A-E, three independent samples of random primed cDNA (biological replicates, N=3) were prepared from strains grown at 30°C and subjected to Real-time PCR analysis using primer sets specific for each mRNA. Transcript copy numbers/2 ng cDNA of each strain were normalized to *SCR1* RNA levels in respective strains and are shown as means ± SE. The statistical significance of difference as reflected in the ranges of p-values estimated from Student’s two-tailed t-tests for a given pair of test strains for each message are presented with following symbols, * <0.05, **<0.005 and ***<0.001, NS, not significant.

### Tif4631p appears to mediate the interaction of CTEXT with the nuclear exosome

To evaluate if the N-terminal RRM1 domain of Tif4631p plays any role in either nucleation of the CTEXT proteome or assists in the interaction with the core nuclear exosome (EXO-11), we purified Tif4631p-ΔRRM1-TAP using the tandem affinity purification as mentioned above and subsequently assess whether the components of CTEXT or core exosome could be detected using western blotting. As shown in Fig. 7A, a comparison of the profile of the co-purifying proteins with a full length Tif4631p-FL and Tif4631p-ΔRRM1 suggest that almost all the components of CTEXT and NNS component Nrd1p was detectable in both cases. In contrast, none of the core exosome components, namely Rrp6p and Rrp4p were not detectable (Fig. 7A). This data is strongly indicating that the N-terminal RRM1 domain plays a vital role in mediating the physical association with the core nuclear exosome. However, further experiments are required to gain an insight into the details of the physical interactions between these protein complexes that play vital functional role in the nuclear mRNA surveillance in *S. cerevisiae*.

**Figure 7.**
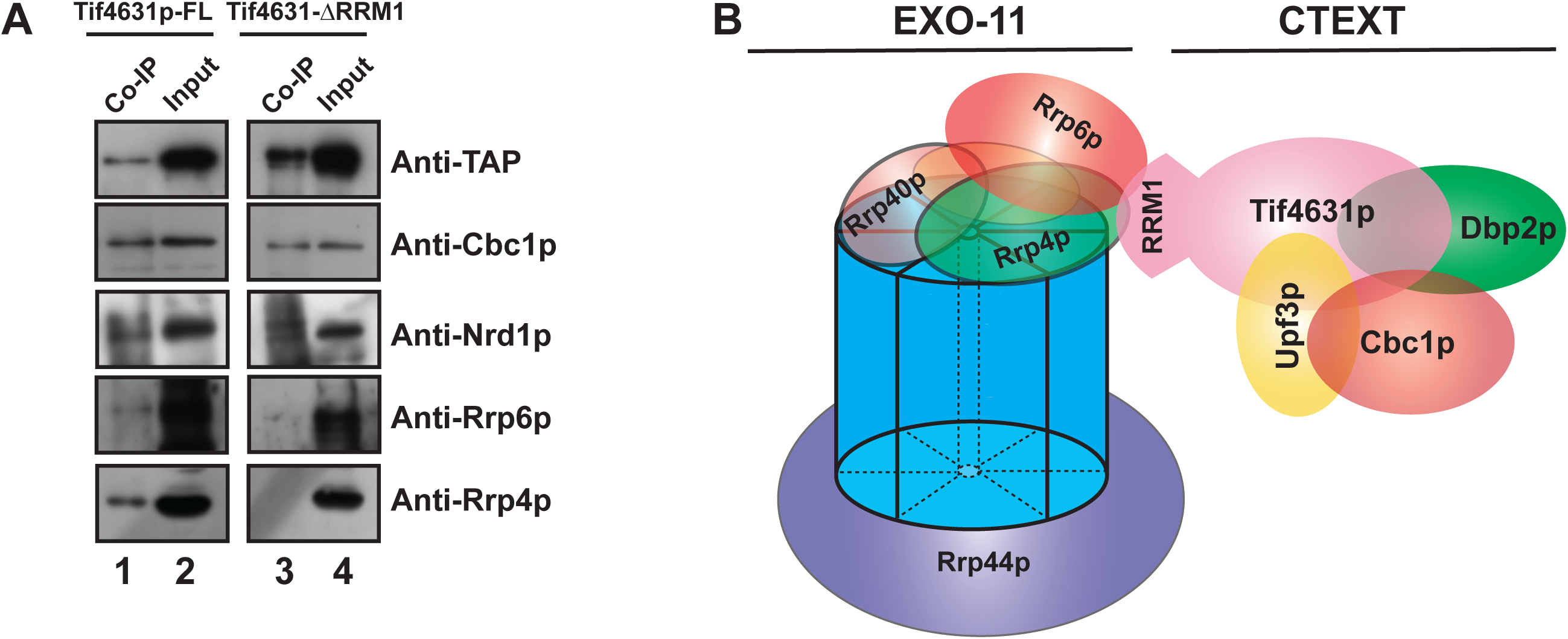
The N-terminal RRM1 domain of Tif4631p specifically mediates the interaction of CTEXT with the core nuclear exosome complex. (A) Result from a representative co-immunoprecipitation experiment using either a TAP-tagged Tif4631p or TAP-tagged Tif4631p-ΔRRM1 as bait. Protein extracts from either a TAP-tagged Tif4631p or TAP-tagged Tif4631p-ΔRRM1 were processed according to the protocol in Fig. 4A, electrophoresed on denaturing SDS gel followed by the detection of the specific CTEXT and exosome components using western blotting by using specific antibodies. The input (Lanes 1 and 4) in case of each sample is 100 μg of whole-cell protein extract from the respective strains, while the Co-IP (Lanes 2 and 3) for each lane represents 20 μl of the eluate in SDS-gel loading buffer from respective strains. (B) Proposed model of interaction of CTEXT with the nuclear exosome. The interaction is likely mediated via the N-terminal RRM1 domain of the Tif4631p as indicated from the Co-IP experiment described in Fig. 7A. The model portrays the interaction of the CTEXT that is mediated by the RRM1 domain of Tif4631p (indicated as a separate domain) with the core nuclear exosome (mediated presumably via Rrp6p and Rrp4p). Other components of CTEXT are coded with different colors and are also indicated.

## Discussion

Control of mRNA abundance via their selective and preferential destruction is a novel mode of regulation of gene expression in eukaryotes. **DRN** (**<u>D</u>**ecay of **<u>R</u>**NA in the **<u>N</u>**ucleus) defines one such mRNA degradation system that, along with the nuclear exosome targets and degrades aberrantly long 3′-extended and export-defective messages in the nucleus (Maity et al. 2016). Moreover, besides targeting the aberrant messages, DRN/exosome also dictates the cellular repertoire of several non-aberrant functional mRNAs, termed ‘special mRNAs’ by their preferential decay in the nucleus. DRN/exosome therefore plays a vital functional role in promoting the generation of functional mRNAs. A comprehensive and structural definition of DRN and how it assists the nuclear exosome, therefore constitutes a major step towards understanding the complex process of nuclear mRNA surveillance.

Previous genetic efforts identified three crucial components of this decay system, Cbc1/2p, Tif4631p, and Upf3p as well as established several classes of aberrant and special RNA substrates, which are targeted by DRN (Das et al. 2000, 2003, 2006, 2014; Maity et al. 2016). However, a convincing evidence of physical interactions among the three previously identified DRN components and their molecular tie with the core nuclear exosome was previously lacking. In this investigation, we used the major DRN component Tif4631p as a bait to co-purify the interacting partners in order to gain an insight into the physical associations among the DRN/core nuclear exosome components. The choice of using this protein was based on the previous evidence where Tif4631p might serve as a nucleating center of various functional protein complexes in different aspects of RNA metabolism (Goyer et al. 1993; Tarun Jr. and Sachs 1995, 1997; Tarun Jr. et al. 1997; Kafasla et al. 2009; Das and Das 2016). Consequently the investigation was initiated with a functional analysis of various known functional domains of Tif4631p to assess which domain is crucial for the nuclear mRNA surveillance. Our analysis revealed that the RRM1 and PAB domain located in the extreme N-terminal end of Tif4631p plays a fundamental role in mRNA surveillance. This conclusion is based on the findings that deletion of either of the N-terminal RRM1 or PAB domains of Tif4631p led to the stabilization of all kinds of aberrant mRNAs tested (Fig. 1 and 2). Subsequent analysis of their expression level showed that although the Tif4631-ΔPAB1 mutant did not display any expression, the Tif4631p-ΔRRM1 mutant exhibited a similar level of expression of full length protein (Fig S2D). This observation refrain us to conclude anything about the Tif4631p-ΔPAB1 mutant protein. Furthermore, the deletion of RRM1 in Tif4631-ΔRRM1 protein domain did not affect its nuclear localization (Fig. S2 E) thereby prompting us to conclude that the N-terminal RRM1 is pivotal for carrying out its function in the nuclear surveillance of 3′-extended and export-incompetent mRNAs.

A careful scrutiny in the N-terminal domain of Tif4631p revealed that this region is highly conserved among the Tif4631p proteins from other species of *Saccharomyces* (Park et al. 2011) that was predicted to be relatively less structured and contains mostly charged residues. The extreme N-terminus of this domain is highly negative that is followed by a positively charged region, which is consistent with the RNA binding activity of this domain (Cawley and Warwicker, 2012). Thirteen invariant residues across different yeast species are present in this region of which eleven of these residues also remain identical in equivalent positions between Tif4631p and Tif4632p isoforms. Since either of the Tif4631p or Tif4632p participates in the nuclear mRNA surveillance (Das et al. 2014), one or more of these invariant residues may predictably constitute the crucial amino acids, which likely plays a crucial role in mediating the interaction between CTEXT-Exosome components, Rrp6/4p. Future experiments directed towards evaluating their functional contribution in nuclear mRNA surveillance in baker’s yeast should throw light on this issue.

Our co-purification procedure using the IgG and Calmodulin columns in tandem following the procedures described by Puig et al (Puig et al. 2001) yielded a large number co-purifying proteins that include the components of CTEXT, the core nuclear exosome (Rrp4p), TRAMP (Mtr4p), NNS (Nrd1p), splicing (Prp proteins), translation (Pab1p, eIF4B, Tif2B), and non-sense mediated decay (Nmd2p, Dcp2p, and Ski2/3p) (Fig. 3D). This observation is perfectly meaningful since Tif4631p is a shuttling protein (McKendrick et al. 2001; Kafasla et al. 2009) and is distributed both in the nucleus and cytoplasm. Notably, Tif4631p was previously identified and functionally characterized as a well-understood cytoplasmic protein which plays a fundamental regulatory role in the initiation of cap-dependent translation (Goyer et al. 1993; Tarun Jr. et al. 1997; Sonenberg and Hinnebusch 2009; Clarkson et al. 2010; Park et al. 2011). More recently both the isoforms of this protein were implicated in the nuclear mRNA splicing of a small group of intron-containing pre-mRNAs (Kafasla et al. 2009) and nuclear mRNA surveillance (Das et al. 2014; Maity et al. 2016). Thus, the presence of so many splicing factors and translation initiation and cytoplasmic non-sense mediated decay factors revealed in our LC-MS/MS results is in perfect agreement with these observations. To corroborate the results of the LC-MS/MS results, we extend the double column purification by further carrying out the co-immunoprecipitation followed by their detection by western blot analysis. This approach led to the detection of the same sets of proteins – previously known CTEXT components, Cbc1p, Upf3p and newly identified factor, Dbp2p, exosome components Rrp4p, and NNS component Nrd1p. Interestingly, while our LC-MS/MS data was unable to detect the exosome component Rrp6p, the Co-immunoprecipitation followed by western blot analysis consistently and reproducibly identified Rrp6p (Fig. 2), the reason which was not clear to us. It is possible that the association of Rrp6p with Tif4631p could be sub-stoichiometric and consequently the low abundance of Rrp6p in the sample somehow went undetected by LC-MS/MS analysis.

Extension of finding obtained from the co-immunoprecipitation experiment using Tif4631p-TAP as a bait led us to conduct additional similar experiments, using any one of the Cbc1p-TAP, Upf3p-HA and Dbp2p-FLAG as bait separately in independent experiments (Fig. 4-5). The data from these experiments validated that same sets of proteins could be detected each time in all of these immunoprecipitation experiments no matter which component was used as bait. Collective data from all these experiments uncovered that Cbc1p, Tif4631p, Upf3p and Dbp2p form a stable complex, which was dubbed CTEXT. Previous findings implicated CTEXT in the nuclear surveillance of the aberrantly long 3′-extended transcripts (additionally requires TRAMP and the exosome) and export defective messages (additionally requires the nuclear exosome) (Maity et al. 2016). Consistent with this observation, our LC-MS/MS analysis detected interaction of Tif4631p with the exosome component Rrp4p and Rrp6p and TRAMP component Mtr4p. Recent studies from our lab also demonstrated the involvement of NNS component, Nrd1p in the recruitment of the exosome component of Rrp6p onto translation elongation-defective and export defective messages to facilitate their rapid degradation (Singh et al. 2020) that validates the observed interaction of CTEXT components with Nrd1p. Taken together, our proteomic data as presented in Figs. 3-5, defines the physical identity and existence of the CTEXT as an independent and integral protein complex that functions as a co-factor of the nuclear exosome to target distinct sets of nuclear mRNAs.

Interestingly, our co-purification procedure also identified Dbp2p (Barta and Iggo 1995; Bond et al. 2001), an ATP dependent DEAD box containing RNA helicase, which qualified as a novel component of CTEXT. Like the other components, Dbp2p also shuttles between the nucleus and cytoplasm and previous studies implicated this factor in the mRNA surveillance and transcription termination (Bond et al. 2001; Cloutier et al. 2012). Like other DEAD box helicases, Dbp2p has a highly conserved helicase core that is involved in ATP binding, hydrolysis and RNA binding and a DEAD box that is characterized by the repeating Asp-Glu-Ala-Asp (DEAD) motif (Linder and Jankowsky 2011). These proteins also display an ATP-dependent RNA helicase activity (Putnam and Jankowsky 2013), which is responsible for unwinding of RNA duplex (Rogers et al. 1999; Yang and Jankowsky 2006) remodeling of RNPs (Tran et al. 2007; Fairman et al. 1999) as well as ATP-dependent fastening of multiprotein complexes (Ballut et al. 2005; Nielsen et al. 2009). Discovery of Dbp2p as the new component of CTEXT is therefore a significant discovery since presumably structural unwinding RNA double strands involving RNP remodeling as well as loading of CTEXT and the exosome complex onto the target RNAs constitute essential activities of an exosomal co-factor. Since an RNA helicase activity associated with the CTEXT (DRN) was undetected till date, this discovery adds another functional dimension to the CTEXT. Consistent with this idea most of the other known exosomal co-factors contain at least one RNA helicases such as Mtr4p in TRAMP (de la Cruz et al. 1998; LaCava et al. 2005; Vanacova et al. 2005), Sen1p of NNS complex (Ursic et al. 1997), Mtl1p of MTREC/NURS in fission yeast *S. pombe* (Egan et al. 2014; Zhou et al. 2015). It is therefore tempting to speculate that Dbp2p likely possesses an ATP-dependent ‘clamping’ activity of multiprotein complexes on the target RNA like other DEAD box RNA helicases (Ballut et al. 2005; Nielsen et al. 2009), which may turn out to be crucial for the recruitment of the core exosome complex on to the target mRNAs. Consistent with this assumption, actual experiments clearly demonstrated that indeed Dbp2p plays a vital role in the degradation of export-defective messages as the levels of these messages were found to undergo significant enhancement in the *dbp2-*Δ yeast strain relative to wild type strain (Fig. 6). Thus, the identification of Dbp2p as a new member of CTEXT complex marks a central and essential discovery that stemmed from the Tif4631p-TAP co-purification experiments. Future work would be directed to evaluate the exact molecular action(s) that Dbp2p exerts as a part of the CTEXT in the nuclear mRNA surveillance.

The observation that none of the exosome subunits, Rrp4p and Rrp6p could be detected as co-purified components in the co-immunoprecipitation experiment involving the TIF4631p-ΔRRM1 protein lacking its N-terminal RRM1 deletion as bait (Fig. 7A) is supportive of the conclusion that the Tif4631p (and its N-terminal RRM1 domain) bridges a physical interaction with the core-exosome complex (Fig. 7A). This association is likely mediated through the Rrp6p and Rrp4p subunits of the core exosome (Fig. 7B). Thus, the abolition of the degradation of special mRNAs and export defective transcripts (Fig. 1 and 2) phenotype associated with the TIF4631p-ΔRRM1 mutant protein could indeed be correlated to the abrogation of the CTEXT-core exosome interaction. This observation therefore validated our original hypothesis that Tif47631p plays a seminal role in mediating the physical interaction between core exosome with the CTEXT. Remarkably, this RRM1 domain (aa 1-82) were previously shown to interact with the PABP interacting PAB1 domain (aa 188-299) while interacting with Poly(A) bound Pab1p during the translation-initiation (Park et al. 2011). This interaction is partially facilitated by the two highly conserved Box1, Box2 and Box3 regions located between the aa 94-111, aa 128-150 and aa 197-224 in the primary structure of Tif4631p (Park et al. 2011) in which Boxes 1 and 2 are acting as a bending arch like structure that brings the RRM1 and Box3/PAB domains in close proximity to each other (Park et al. 2011). Interestingly, these same regions from the two isoforms of Tif4631p and 32p are also highly conserved with the invariant amino acid residues. It remains to see if these Boxes 1, 2 and 3 of Tif4631p impart a similar functional role in mediating the physical association with the core nuclear exosome complex via Rrp6p and Rrp4p. Future research directed towards the functional analysis of these highly conserved regions will further unfold their functional role in the nuclear mRNA surveillance in baker’s yeast.

## Data Availability

All the raw data leading to the establishment of the final results presented here are available in the supplementary material as a separate excel file ‘**SahaUDasS21 NAR Res Art Raw Data**’.

## Acknowledgments

We gratefully acknowledge Dr. Scott Butler (Department of Microbiology and Immunology, University of Rochester, Rochester, NY, USA) for kindly sharing the antibodies against Cbc1p and Rrp6p, Dr. Stephen Buratowski (Department of Biological Chemistry and Molecular Pharmacology, Harvard Medical School, Boston, MA, USA) for kindly sharing the antibody against Nrd1p and Dr. Elizabeth Tran (Department of Biochemistry, Purdue University, West Lafayette, IN, USA) for kindly sharing isogenic wild-type and *dbp2-*Δ mutant strains as gifts. The TAP-plasmid was a kind gift from Dr. Sander Granneman (University of Edinburgh, United Kingdom). We thank members of the Das Laboratory for critically reading this manuscript. We also thank the anonymous reviewers for their critical comments and constructive suggestions, which greatly enriched this manuscript.

## Abbreviations

CTEXT: **<u>C</u>**BC-**<u>T</u>**if4631p-dependent **<u>EX</u>**osome **<u>T</u>**argeting
TRAMP: **<u>TR</u>**f4p/5p-**<u>A</u>**ir1p/2p-**<u>M</u>**tr4p-**<u>P</u>**olyadenylation,
DRN: **<u>D</u>**ecay of m**<u>R</u>**NA in the **<u>N</u>**ucleus;
NLS: **<u>N</u>**uclear **<u>L</u>**ocalization **<u>S</u>**ignal
NES: **<u>N</u>**uclear **<u>E</u>**xport **<u>S</u>**ignal

