## Supplementary figures and images for "The CTEXT complex in *Saccharomyces cerevisiae* plays a crucial role in degrading distinct sets of aberrant mRNAs by the nuclear exosome"

### Supplementary Figure S1

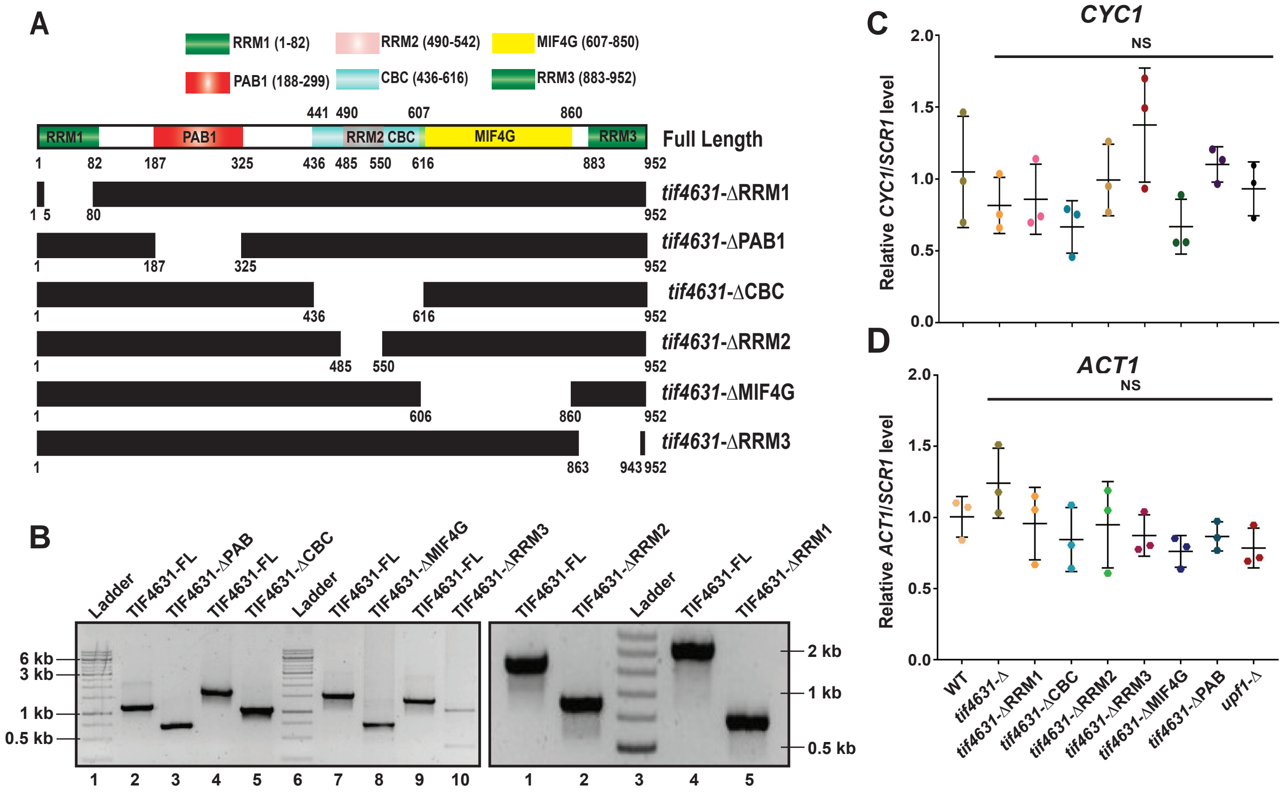


**Figure S1**

Saha et al. (2021)

SahaU21 NAR Res Art Supplementary Fig. S1.pdf

### Supplementary Figure S2

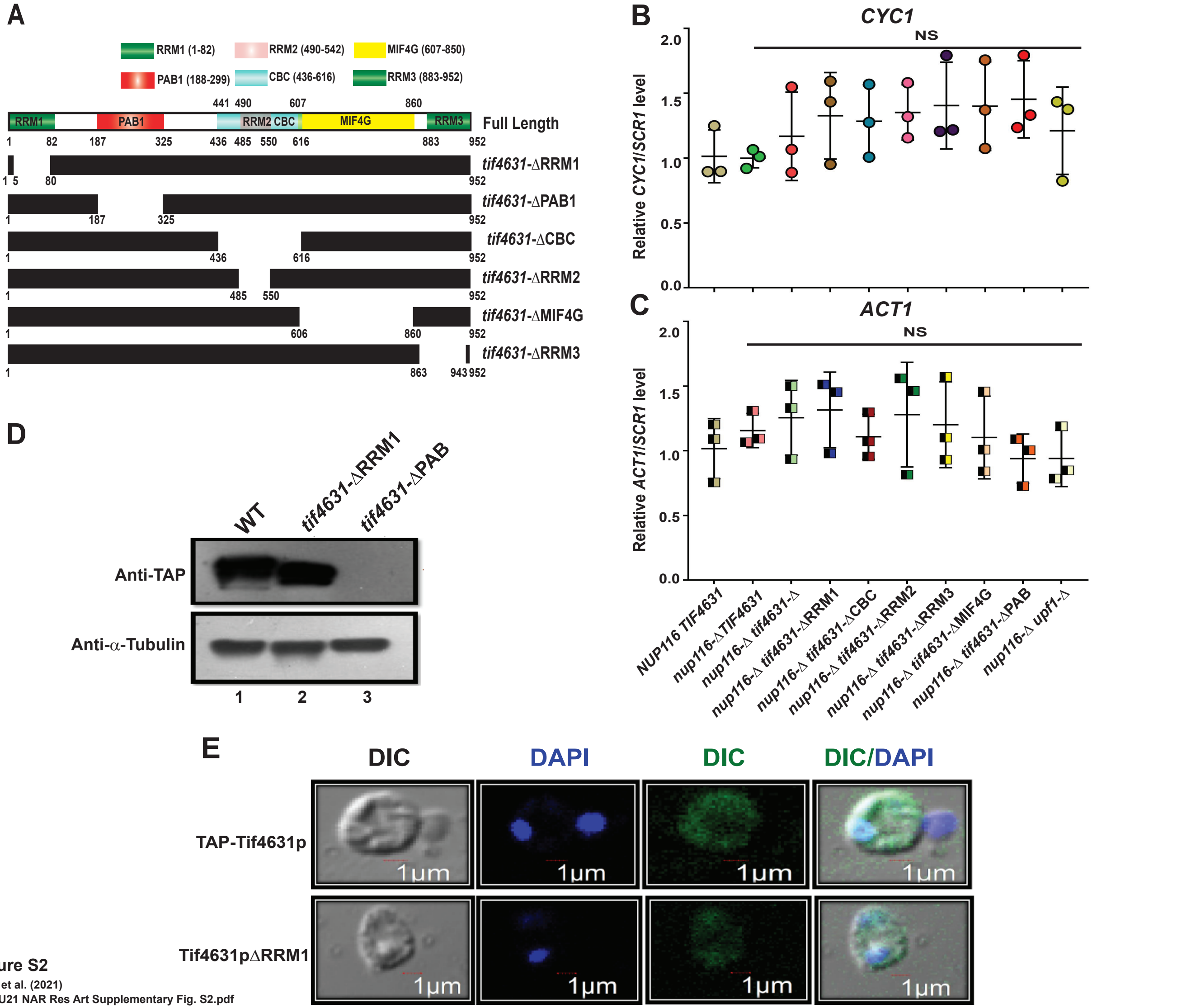


**Figure S2**

Saha et al. (2021)

SahaU21 NAR Res Art Supplementary Fig. S2.pdf
